# FYN tyrosine kinase, a downstream target of receptor tyrosine kinases, modulates anti-glioma immune responses

**DOI:** 10.1101/608505

**Authors:** Andrea Comba, Patrick J Dunn, Anna E Argento, Padma Kadiyala, Maria Ventosa, Priti Patel, Daniel B Zamler, Felipe J Nunez, Lili Zhao, Maria G Castro, Pedro R. Lowenstein

## Abstract

**Background:** High grade gliomas are aggressive and immunosuppressive brain tumors. Molecular mechanisms that regulate the inhibitory immune tumor microenvironment (TME) and glioma progression remain poorly understood. FYN tyrosine kinase is a downstream target of the oncogenic receptor tyrosine kinases pathway and is overexpressed in human gliomas. FYN’s role *in vivo* in glioma growth remains unknown. We investigated whether FYN regulates glioma initiation, growth and invasion.

**Methods:** We evaluated the role of FYN using genetically engineered mouse glioma models (GEMM). We also generated FYN knockdown stem cells to induce gliomas in immune-competent and immune-deficient mice (NSG, CD8−/−, CD4−/−). We analyzed molecular mechanism by RNA-Seq and bioinformatics analysis. Flow cytometry was used to characterize immune cellular infiltrates in the FYN knockdown glioma TME.

**Results:** We demonstrate that FYN knockdown in diverse immune-competent GEMMs of glioma reduced tumor progression and significantly increased survival. Gene ontologies (GOs) analysis of differentially expressed genes in wild type vs. FYN knockdown gliomas showed enrichment of GOs related to immune reactivity. However, in NSG, CD8−/− and CD4−/− immune-deficient mice, FYN knockdown gliomas failed to show differences in survival. These data suggest that the expression of FYN in glioma cells reduces anti-glioma immune activation. Examination of glioma immune infiltrates by flow-cytometry displayed reduction in the amount and activity of immune suppressive myeloid derived cells (MDSCs) in the FYN glioma TME.

**Conclusions:** Gliomas employ FYN mediated mechanisms to enhance immune-suppression and promote tumor progression. We propose that FYN inhibition within glioma cells could improve the efficacy of anti-glioma immunotherapies.

**Key points:** Inhibition of FYN tyrosine kinase in genetically engineered mouse glioma models delays tumor initiation and progression. The oncogenic effects of FYN in vivo are mediated by downregulation of anti-glioma immunity.

**Importance of the Study:** FYN is an effector of receptor tyrosine kinases (RTK) signaling in glioma. However, its role *in vivo* remains unknown. Our study demonstrates that FYN tyrosine kinase is a novel regulator of the anti-glioma immune response. We show that FYN inactivation suppresses glioma growth, increases survival, and enhances anti-tumor immune reactivity. Our findings suggest that suppressing the expression of FYN in glioma cells could provide a novel therapeutic target.

## INTRODUCTION

Glioblastoma multiforme, or high grade glioma (HGG) is the most frequent and aggressive primary tumor of the central nervous system. It is characterized by extensive infiltrative growth and resistance to therapy^1^. Mutated/activated driver genes such as the receptor tyrosine kinases (RTK: EGFR, PDGFR, HGF/MET), tumor suppressor genes (TP53, PTEN/NF1) and downstream RAS/MEK/ERK or PIK3/AKT pathways contribute to the malignity of glioma^2^. FYN, a non-RTK member of the SRC family kinase (SFK), is a downstream proto-oncogene target of the RTK pathway,^3–6^. However, the specific mechanisms by which FYN stimulates glioma growth and invasion remains unknown. FYN is rarely mutated^7^ in human HGG, but is significantly overexpressed.

Several *in-vitro* studies showed that FYN knockdown is associated with decreased cell migration and proliferation of glioma cells^3,8,9^ Nevertheless, *in vivo* human glioma xenograft models of FYN Knockdown in immune-suppressed animals failed to show differences in survival^8^. Therefore, an immune-competent mouse model that enables study of gliomas with FYN knockdown was established.

In this study we demonstrate that FYN, a downstream target of receptor tyrosine kinases signaling, inhibits the anti-glioma immune response. We demonstrated that FYN tyrosine kinase promotes glioma initiation and growth utilizing both immune-competent and immune-deficient mouse glioma models. We observed that GEMM of gliomas and implantable gliomas, both with FYN knockdown displayed an extended survival compared to wildtype FYN tumor bearing immune-competent mice. However, FYN knockdown tumors implanted into immune-deficient mice did not extend survival when compared to controls. Molecular analysis revealed a significant upregulation over-representation of immune related gene ontologies (GOs) in shFYN tumors. The over-representation of immune-related GOs suggest that possibility that FYN is somehow suppressing immune function. Our data suggest that FYN expression in glioma cells suppresses the immune responses by stimulating the expansion and activity of MDSCs in glioma tumor microenvironment. Gliomas have been demonstrated to employ a variety of immunosuppressive mechanism which promotes tumor progression, thus reducing the effectiveness of immunotherapies^10,11^.

Our data uncover a new paradigm of how FYN tyrosine kinase expressed within the tumor cells regulates anti-glioma immune responses. We propose that tumor cell specific inhibition of FYN tyrosine kinase will increase the sensitivity of gliomas to immune attack, and represents a potential target for future treatments of glioma patients.

## MATERIALS AND METHODS

### Glioma cells

Mouse neurospheres cells were derived from genetically engineered mouse models (GEMM) of gliomas. These cells were generated in our lab using the Sleeping Beauty (SB) transposon system^12,13^ All cells were cultured as described in Supplementary data.

### Generation of stable cell lines with FYN knockdown

NP and NPA neurospheres were used to generate stable cell lines with FYN knockdown. The pLenti pLKO-non-target shRNA control vector (SHC002) and two different pLenti-mouse shRNA vectors for FYN were selected from Sigma Aldrich MISSION^®^ shRNA Vectors. The FYN shRNA identification numbers are: TRCN0000023383 (shFYN #1) and TRCN0000361213 (shFYN #2). Cells were infected with the lentivirus as described previously by us^14^. Immunoblotting was used to confirm FYN knockdown. FYN shFYN #2 cells were selected for *in vivo* experiments. Moreover, to validate the specificity of the shFYN and discard any potential off-target effects we performed a rescue experiments as described in detail in Supplementary data.

### Intracranial implantable syngeneic mouse glioma model

Studies were conducted according to the guidelines approved by the Institutional Animal Care (IACUC) and Use Committee at the University of Michigan (approved protocol, PRO00007666 for C57BL/6 immune-competent mice and PRO00007669 for immune-suppressive mice). 3.0 x 10^4^ neurospheres were implanted into the striatum of mouse brains to generate tumors. See supplementary data.

### Genetically engineered mouse glioma model (GEMM) generation for FYN Knockdown

The animal model studies were conducted in C57BL/6 mice (Jackson Laboratory), according to IACUC approved protocol PRO00007617. A FYN knockdown glioma murine model and the appropriate controls were created by the Sleeping Beauty (SB) transposon system as described^12,13^ The genotypes of SB generated tumors were: (i) shp53 and NRAS **(NP)**, (ii) shp53, NRAS and shFYN **(NPF)**, (iii) shp53, NRAS and shATRX **(NPA)**, (iv) shp53, NRAS, shATRX and shFYN **(NPAF)**, (v) shp53, NRAS and PDGFβ **(NPD)**, (vi) shp53, NRAS, PDGFβ and shFYN **(NPDF)**. Design and cloning of shFYN vector is in Supplementary data.

### Immunoblotting

Glioma cells (1.0 x 10^6^ cells) were seeded in a 100 mm dish and grown at various time points as shown in supplementary data.

### Immunohistochemistry of paraffin embedded brains (IHC-DAB)

Immunohistochemistry assay was performed on paraffin embedded tissue as described previously^15^.

### Immunofluorescence of paraffin embedded brains

Brains that were fixed in 4% paraformaldehyde were processed, embedded in paraffin and sectioned as described previously^15^. FYN antibody was conjugated with Alexa Fluor™ 488 Tyramide SuperBoost™ Kit - Goat Anti-Rabbit IgG (# B40922) following the manufacture instructions (Invitrogen-Thermo Fisher Scientific).

### RNA isolation and RNA-Sequencing

SB NP, NPF, NPA, NPAF, NPD and NPDF tumors were studied by RNA-Seq analysis. RNA was isolated using the RNeasy Plus Mini Kit (© QIAGEN) following the manufacture instructions. RNA-sequencing was performed at the University of Michigan DNA Sequencing Core. Detailed analysis is described in Supplementary data.

### Flow Cytometry

For flow cytometry analysis of immune cells within the TME, we generated tumors by intracranial implantation of 3.0 x 10^4^ NPA-NT and NPA-shFYN cells in C57Bl6 mice. Protocol was performed as described before^16^ and detailed in Supplementary data.

### *In vitro* MDSC migration assay

We generated mouse bone marrow-derived MDSC as described by Maringo *et al (**REF**).* We analyzed *in vitro* MDSC (M-MDSC and PMN-MDSC) migration using Transwell^®^ polycarbonate membrane inserts (Corning Inc.) of 6.5 mm diameter and 8 um pore size. Detailed methodology is described in Supplementary data.

### T Cell Proliferation Assays

We measured MDSC immune suppressive activity using the in vitro T cell proliferation assay. MDSCs were purified from TME of NPA-NT and NPA-shFYN tumors from moribund mice. MDSC were purified by flow sorting as Gr-1^high^ (PMN-MDSC) and Gr 1^low^(M-MDSCs) as described^16^. See detailed methodology in Supplementary data.

### Statistical Analysis

All experiments were performed in at least three or more independent biological replicates, depending on the specific analysis. Data is presented as the mean ± SEM. All statistical tests used are indicated within the figure legends and in Supplementary data.

## RESULTS

### FYN was identified as a potential regulator of glioma progression

FYN tyrosine kinase is activated by several receptor tyrosine kinases (RTK), such as EGFR, PDGFR and c-MET, commonly mutated genes in gliomas. Following FYN activation, there are several downstream RAS dependent and RAS-independent signaling pathways such as RAS/MEK/ERK, PIK3/AKT, FAK, PXN, B-catenin, STAT3, SHC and VAV2 leading to changes in proliferation, migration, invasion and cell-cell adhesion **(Fig 1A).**

**Figure 1.**
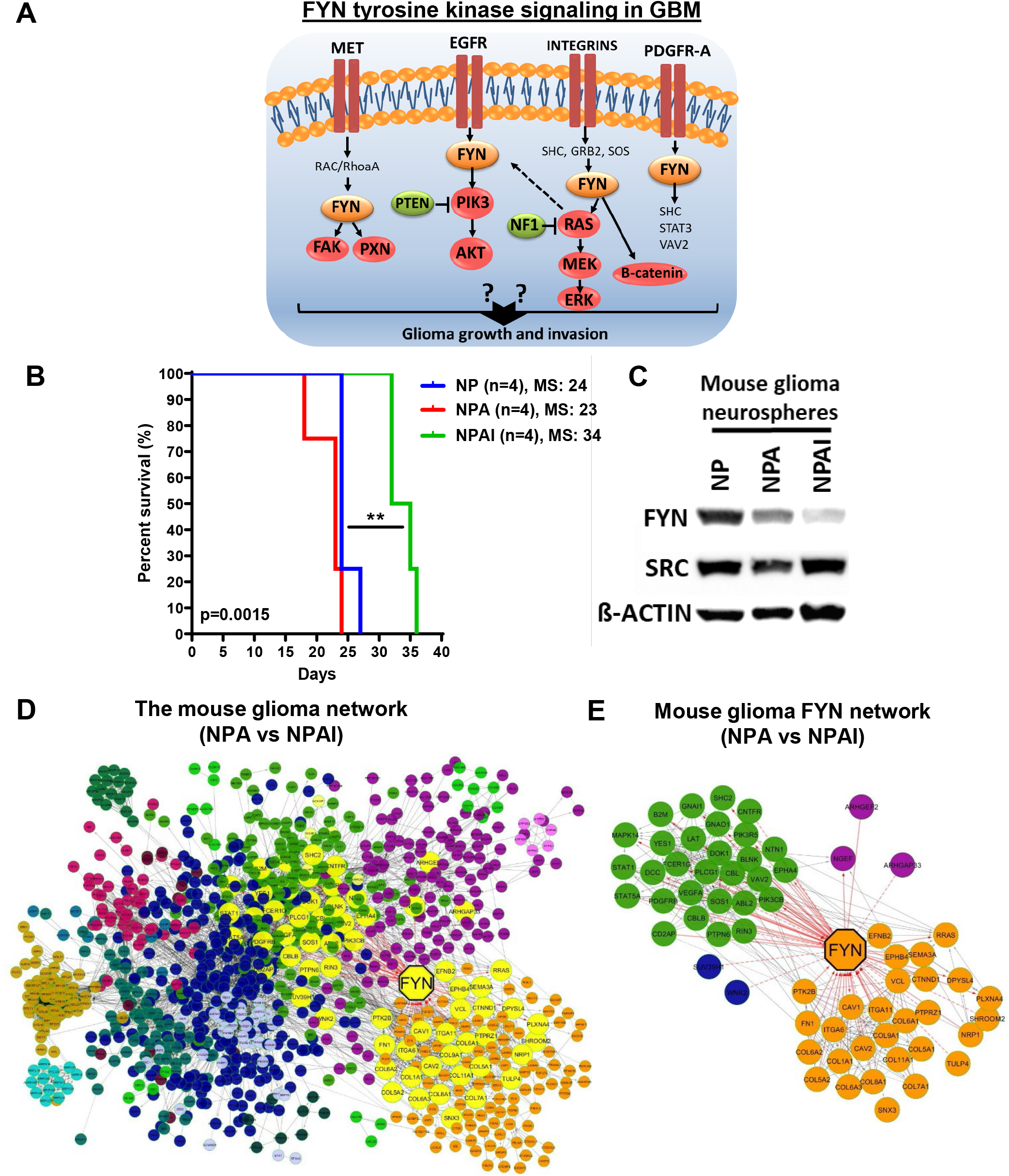
FYN is a potential regulator of aggressiveness in mouse gliomas. **(A)** FYN tyrosine signals downstream from mutated membrane RTK and Integrin driver genes. Red: mutation/amplification, green: mutation/deletion, orange: FYN overexpression. **(B)** Kaplan–Meier survival curves of implantable mouse glioma models shows difference in tumor malignancy. NPAI display an increased survival compared with NP and NPA glioma bearing mice. NPAI (MS: 34 days; n: 4), NP (MS: 24 days; n: 4), NPA (MS: 23 days; n: 4). **(C)** FYN levels correlate positively with glioma cell malignancy in WB analysis comparing FYN and SRC levels in mouse (NP, NPA, NPAI) glioma cells; loading control = β-actin. **(D)** Network of DE genes in high malignancy NPA vs low malignancy NPAI mouse glioma neurospheres. FYN is the largest yellow octagon. Cluster of nodes of identical color represents a module of highly interacting genes. The FYN network is highlighted in yellow. **(E)** Right panel: the FYN network; FYN has a degree of 63. Red lines indicate edges connecting nodes to FYN.

We analyzed RNA-Sequencing and Microarray human data from the Gliovis (http://gliovis.bioinfo.cnio.es) database. According to Rembrandt, TCGA, and Gravendeel databases, FYN mRNA expression levels were higher in different types of human gliomas when compared to normal brain tissue **(Supplementary Fig. S1A)**. We observed that FYN expression was positively correlated with mouse glioma cells aggressiveness. Figure 1B shows that the survival of animals implanted with NP and NPA glioma cells is significantly shorter than that of animals implanted with NPAI cells. In accordance, Western blot analysis in Figure 1C, demonstrates that the levels of FYN, but not SRC, are higher in NP and NPA cells compared to NPAI cells.

To further analyze the importance of FYN in glioma malignancy, we investigated differential expression (DE) of genes in highly aggressive glioma NPA-neurospheres (NRAS, shp53, shATRx, IDH-Wild-type) compared to NPAI-neurospheres (NRAS, shp53, shATRx, IDH1^R132H^), of lower aggressiveness^13^. The network of DE genes identified FYN to be one of the most highly connected nodes (Degree: 63; 4^th^ node from top node), a hub in the network **(Fig. 1C-D). Fig. (1D)** and **(1E)** display the network of FYN, a set of nodes directly connected to FYN. Higher magnification of the networks is shown in **Supplementary Figure S1A-B.** Functional network GO term analysis disclose Cell Proliferation, Cell migration, MAPK cascade, Positive regulation of PI3K signaling, VEGF receptor signaling, Cellular response to PDGF as significant GO Biological Processes involving FYN **(Supplementary Fig. 1B)**. The list of GO terms with their respectively q and p values are shown in **Supplementary, Table S1.**

### Loss of FYN reduces tumor malignancy and prolongs survival of mice harboring GEMM tumors

The Sleeping Beauty Transposon System GBM model (GEMM) was used to understand the function of FYN^12^. We generated a FYN-deficient genetically engineered mouse glioma model (**Supplementary Fig. S3A and B**). We corroborated the efficacy of the shRNAs of FYN by WB analysis (**Supplementary Fig. S3C**). The shFYN-(b) was selected for the GEMM glioma generation. We generated tumors harboring wildtype or FYN knockdown in the presence of various genotype combinations (**Supplementary Fig. S3D**).

Downregulation of FYN increased median survival (MS) in comparison to wildtype FYN control groups (**Fig. 2A, B and C**) in all genotype models. NPF group displayed an increased MS of 131 days compared with 94 days in the NP control group (**Fig. 2A**). The experimental group with knockdown of FYN in the context of ATRX loss (NPAF) exhibited an increased survival (MS: 142 days) compared with the NPA control (MS: 80 days), (**Fig. 2B**). In the third experimental group, FYN knockdown plus PDGFβ ligand upregulation (NPDF), also displayed an increased MS of 108 days compared with the NPD control group (MS: 69 days) (**Fig. 2C**). We corroborated the downregulation of FYN protein in all experimental groups as shown in **Fig. 2D, 2E** and **2F** respectively. Histopathology analysis of tumors showed no evidence of significant differences in glioma malignant pathological markers (**Fig. 2G, H, I and J**). We further evaluated cellular proliferation of the tumor. Quantification demonstrated a significant decrease in the ratio of P-H3-S10 cells per total cells in FYN knockdown groups (**Fig. 2K and I**). These data demonstrate that FYN downregulation increases animal survival by decreasing tumor initiation, development and proliferation.

**Figure 2.**
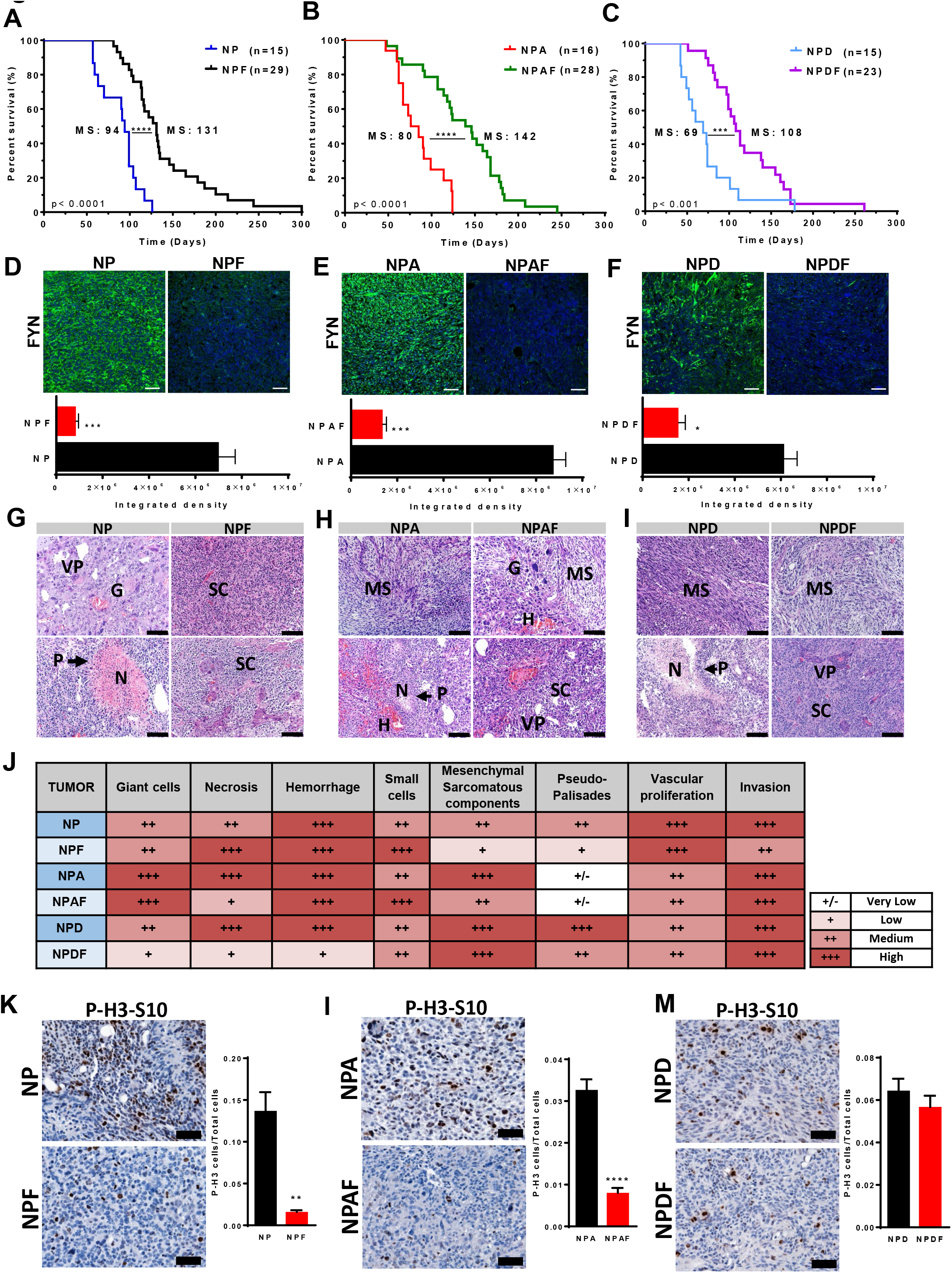
Knocking down FYN in GEMM prolongs animal survival. **(A-C)** Kaplan–Meier survival curves of SB mouse glioma models demonstrate that animals bearing FYN knockdown tumors have increased median survival (MS). **(A)** NP (MS: 94 days; n: 15) versus NPF (MS: 131 days; n: 29). **(B)** NPA (MS: 80 days; n: 16) versus NPAF (MS: 142 days; n: 28). **(C)** NPD (MS: 69 days; n: 15) versus NPDF (MS: 108 days; n: 23). Log-rank (Mantel-Cox) test; *** p<0.001, ****p<0.0001. **(D-F, top)** FYN expression in tumors (green = tumor; and Blue = DAPI stained nuclei), quantified as fluorescence integrated density using Image-J **(D-F, bottom)**; n=5 per condition, scale bar = 50 μm. Ten random fields per tumor section per animal were imaged. Bars ±SEM are shown; (***p<0.001, *p<0.05 using linear mixed effect models) (**G-I)**. Histopathological analysis was performed in tumor sections stained with H&E; shFYN tumors were compared with controls. Scale bars: 100 μm. P: pseudo-palisades, N: necrosis, H: hemorrhage, VP: vascular proliferation, MS: mesenchymal component, SC: small cells, G: giant cells. **(J)** Table representing histopathological semi-quantitative analysis: very low (+/-), low (+), medium (++) and high (+++). **(K-M)** Cell proliferation analysis: Positive P-H3-S10 cells were counted by Image-J software. Scale bars: 50 μm. P-H3-S10 positive cells per total cells in the visual field; n=5. Ten fields of each section were selected at random. Error bars represent ±SEM; linear mixed effect models, ***p<0.001, **p<0.01.

### RNA-Seq and bioinformatics analysis reveals increased representation of immune ontologies in FYN knockdown glioma models

RNA-sequencing and bioinformatics analysis was used to discover changes in gene ontologies (GOs) that could help us understand the mechanism by which FYN knockdown leads to the inhibition of tumor growth and progression. Genomic studies were performed in the following GEMM groups: NPF vs NP, NPAF vs NPA, and NPDF vs NPD. RNA-Seq analysis revealed a group of 515 DE genes in NPF vs NP (205 upregulated genes and 310 downregulated genes; 1295 DE genes in NPAF vs NPA (469 upregulated and 826 downregulated) and 630 DE genes in NPDF vs NPD (565 upregulated and 65 downregulated) (**Supplementary Fig. S4A-D**). Using network analysis (Cytoscape) we analyzed the functional interaction of the DE genes resulting from FYN knockdown (**Supplementary Fig. S4**). The analysis of the network interactions revealed several genes which represent hubs and thus potential regulators of the network functions (**Supplementary Fig. S5 A-C**). We found in NPF vs NP that STAT1, ITGA2, ITGA3, ITGA9, GNA14, CAMK2A represent highly connected hubs of the network. In NPAF vs NPA the most connected genes on the network were NFKB1, STAT1, SHC1, ITGB2, FN1, VCL, ITGB7, CAMK2A. In NPDF vs NPD the hubs of the network are represented by FYN, STAT1, SYK, RAC2, VAV1, PIK3CG, ITGB2. These genes play an essential role in gene regulation and biological processes. Moreover, the functional network analysis found the following pathways as commonly overrepresented in all three shFYN tumors independent of the genetic background of the tumors: Extracellular matrix organization, Focal adhesion, IL12-mediated signaling events, Integrin signaling pathway, Pathways in cancer, PI3K-Akt signaling pathway and Cytokine-cytokine receptor interaction (**Fig. Supplementary S6A-C and Supplementary Table S2A-C S3**). As is shown in the figures, other pathways were specifically impacted for each genetic condition. Further, GOs analysis performed by I-PathwayGuide platform (Advaita Corporation, MI, USA), corrected for multiple comparisons using Elim pruning method, is compatible with the hypothesis that FYN knockdown mediates an activation of immune response among all GEMM of glioma. The analysis shows that the set of top over-represented GO terms, in shFYN glioma for each individual genetic background, are mostly related to immune functions **(Figure 3 A-D and Supplementary table S3 A-C).** Furthermore, to identify common Gene ontologies (GO) shared between all GEMM glioma models for FYN downregulation (NPF vs NP, NPAF vs NPA and NPDF vs NPD) DE genes of each genetic background were compared by meta-analysis. We encountered 58 common over-represented GO terms shared by all three FYN knockdown glioma models **(Figure 3A).** Significantly over-represented GOs in the meta-analysis were associated with immune biological functions including “cellular response to interferon-gamma”, “cellular response to interferon-beta”, “antigen processing and presentation of exogenous peptide antigen via MHC class II”, “cellular response to interleukin-1”, “myeloid dendritic cell differentiation”, “positive regulation of T cell proliferation” between others **(Figure 3E)**. These signaling pathways and GO terms represent potential mechanism by which FYN knockdown decreases glioma malignancy in NPF, NPAF and NPDF glioma models. Details of the selected GO terms are shown in **Supplementary Table S4A-C**.

**Figure 3.**
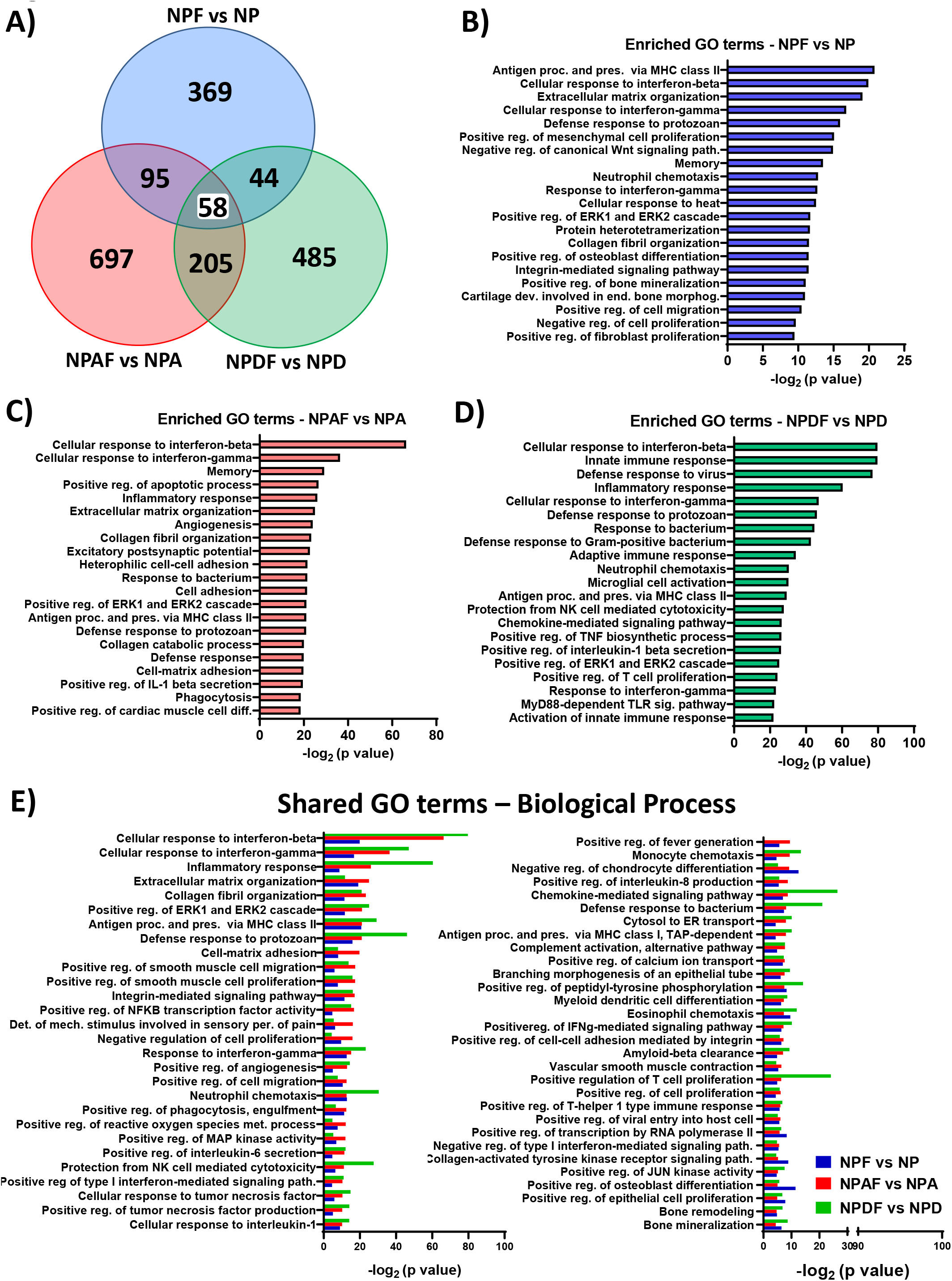
Enrichment in immune-related pathways as potential mechanisms involved in FYN-mediated tumor growth control. Overrepresented Gene Ontology (GO) biological term in the DE genes dataset was carried out using iPathwayGuide algorisms. Over-representation approach to compute the statistical significance was used. p-value is computed using the hypergeometric distribution and corrected for multiple comparisons using *elim* pruning methods. Minimum DE genes/term of 6. **(A)** Venn diagram of GOs Biological Process. Meta-analysis of different genetic glioma models shows individual GO terms and 58 common biological process shared between all groups. For GO analysis, genes with 0.05 p-value and a log fold change of at least east 0.585 absolute value were considered significant. **(B, C, D)** Graph representing the top 20 most significant enriched GO terms identified for each individual genetic glioma model. **(B)** Enriched GO terms of NPF vs NP. **(C)** Enriched GO terms of NPAF vs NPA. **(D)** Enriched GO terms of NPDF vs NPD. **(E)** Bars graph showing the significant over-represented 58 common GO terms biological process identified by meta-analysis.

### The role of FYN in glioma growth, malignancy, and immune response interaction

To test the hypothesis that FYN depletion stimulates anti-tumor immune responses *in vivo,* we implanted FYN knockdown cells in immune-competent and immune-deficient mice. Preceding the *in vivo* experiments We identified by Western blot that FYN was successfully downregulated. Both shRNAs were specific for FYN and does not have effect on SRC tyrosine kinase. Decreased expression of SFK phosphorylation site P-Y416 and P-Y530 were observed (**Supplementary Fig. S7A and B**).

To test if potential off-target effects due to sh-FYN influence our results, we performed an *in vitro* rescue experiment. NPA-shFYn cells transduced with a plv-cherry-FYN vector expressing a silent-mutated form of FYN gene, designed to be resistant to the shRNA of FYN, were used to rescue the shFYN induced knockdown. Expression of this construct counteracted the inhibition of sh-FYN (**Supplementary Fig. S7C**). We then tested the biological effects of FYN expression on cell viability. We observed a decreased proliferation in NPA-shFYN cells compared to NPA-NT. FYN overexpression within NPA-shFYN cells reversed the effects of FYN knockdown (**Supplementary Fig. S7D**). The *in vivo* study showed that inactivation of FYN in glioma cells strongly suppress tumor growth and increase survival in immune-competent mice (C57BL/6) **(Fig. 4A-B)**, but the effects in immune-compromised (NSG) mice was negligible (**Fig. 4C-D**). FYN knockdown group displayed a MS of 34 days compared to the NP-NT control group (MS: 27 days) (**Fig. 4A Fig 5A**). Also, NPA-shFYN had a significantly higher median survival (MS: 30 days) than the NPA-NT control group (MS: 20) (**Fig. 4B**). Moreover, *in vivo* bioluminescence analysis of the tumors at 13 dpi showed that NP-FYN knockdown tumors exhibited lower signal. Tumor size evaluation at necropsy showed a correlation with *in vivo* tumor bioluminescence analysis (**Supplementary Fig. 7E and F**). These results validate the role of FYN in immune-competent mice. However, tumor induction in NSG immune-deficient mice with NP-shFYN tumors did not exhibited a significant difference in survival when compared to the control NP-wtFYN tumors (MS: 24 vs 26 dpi) **(Fig. 4C).** In mice bearing NPA-shFYN tumors a minor, yet statistically significant increase in MS (22 vs 24 dpi) was observed (**Fig. 4D**). Moreover, implantation of NP-shFYN and NPA-shFYN tumors in CD8 Knockout mice showed no difference in survival compared to the controls tumor bearing mice **(Fig. 4E and Supplementary Fig. S7G).** Implantation of NPA-shFYN tumors in CD4 Knockout mice showed no difference in survival (**Fig. 4F**). Collectively, these results demonstrate that enhanced survival in FYN knockdown tumors is mediated by the immune system, including CD8 and CD4 T cells.

**Figure 4.**
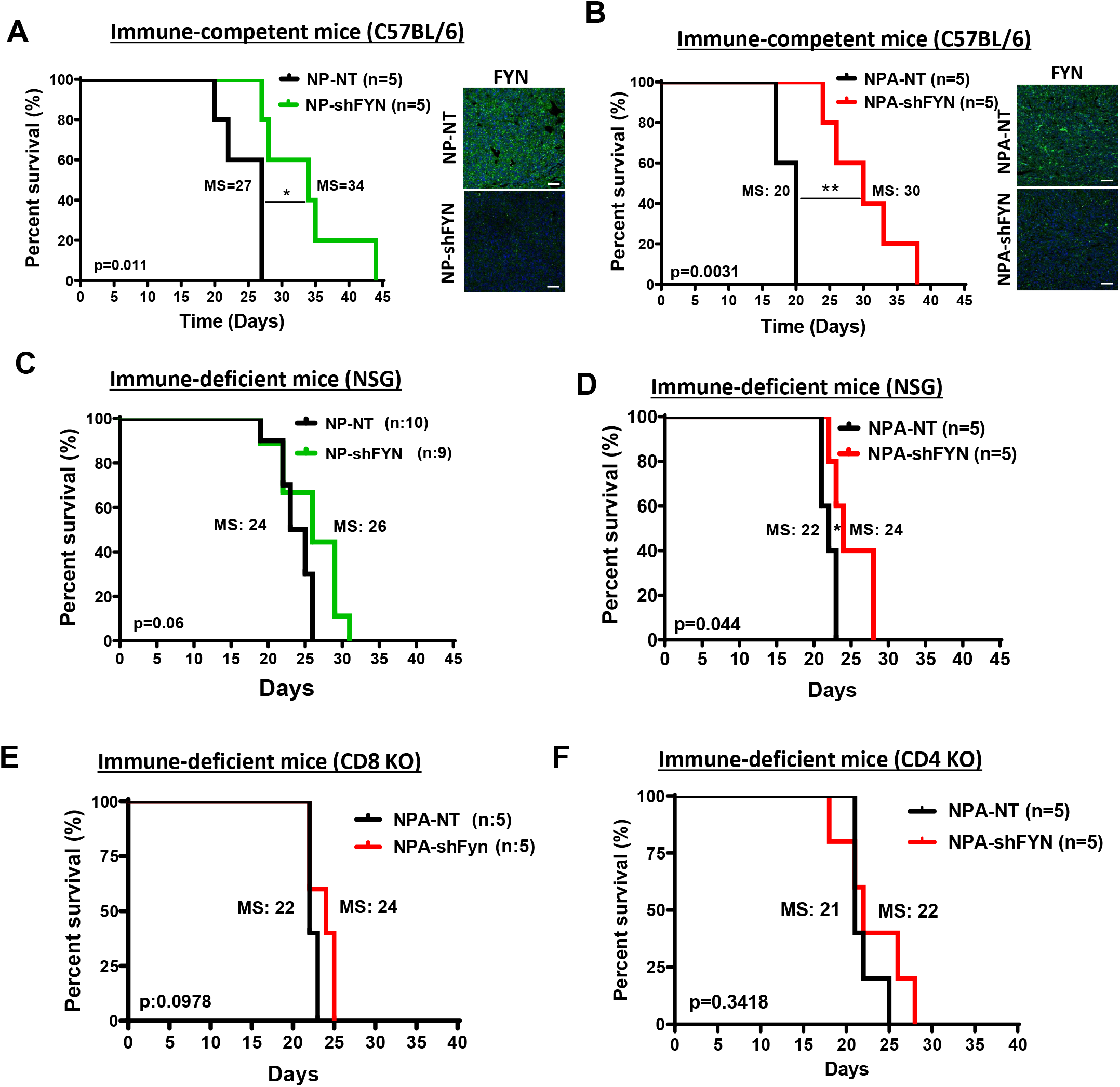
Tumor growth delay and increased survival in FYN downregulated gliomas is immune mediated. **(A-B)** Kaplan–Meier survival curves for glioma in C57BL/6 immune-competent mice. **(A)** NP-FYN knockdown gliomas displayed significant increases in MS: 27 vs 34 days; *p=0.011. **(B)** Mice bearing gliomas with NPA-FYN knockdown displayed significant increase in MS: 20 vs 30 days; **p=0.0031 compared to the control; n=5. FYN expression was detected by immunofluorescence. Tumor: green; Nuclei: blue; scale bars: 50 μm. **(C-D)** Kaplan–Meier survival curves in NSG immune-compromised mice. **(C)** NP-FYN knockdown gliomas displayed no significant increases in MS (24 vs 26 days; p=0.06) n=9/10. **(D)** NPA-FYN knockdown displayed a minor increase in MS (22 vs 24 days; *p=0.044); n=5. **(E-F)** Kaplan–Meier survival curve for NPA-NT vs NPA-shFYN glioma in **(E)** CD8 and **(F)** CD4 KO immune-deficient mice. No significant difference was observed in survival. For each implantable model n=5 was used. Statistics were assessed using the log-rank Mantel-Cox test.

As was observed above the survival benefit of sh-FYN is larger in GEMM than in implantable tumors. The reason for the different survival benefit is that in the GEMMs, tumors originate de novo as a result of the genetic modification of neural stem cell progenitors in one day old pups. In the implantable glioma model, tumors are induced by intracranial implantation of 30,000 cells. Median survival is 80, 94 days in GEMM, and 27, 20 days in implantable tumors. Percentagewise, sh-FYN increases survival by 139%, 177% in GEMM (NP, NPA), and by 125%, 150% in NP, NPA implantable tumors (**Fig 2A-B**; **Fig. 4A-B**). We believe that the survival benefit of shFYN is reduced in implantable gliomas due to their increased growth rate.

### FYN downregulation in glioma reduces Myeloid derived suppressive cells (MDSC) amount and activation markers, and inhibitory potency within the tumor microenvironment (TME)

To determine whether downregulation of FYN in gliomas has an effect on the immune response, we examined the immune cellular infiltrates in the TME. First, we analyzed the role of T cells in the TME of shFYN tumors. No significant difference in the frequency of CD4^+^ T cells (CD45^+^, CD3^+^, CD4^+^) or CD8^+^ T cells (CD45^+^, CD3^+^, CD8^+^) were observed in NPA-NT versus NPA-shFYN tumors (**Fig. 6A, B, C and D).** However, we found a reduction in CD8 T cells that express PD1, a marker of T cell exhaustion **(Fig. 5E-F).** In gliomas, increases in MDSCs are an important mechanism of anti-tumor immune evasion^16^. Therefore, we evaluated the expansion of MDSCs mediated immunosuppression in glioma TME. MDSCs were identified as monocytic M-MDSC (CD45^+^, CD11b^+^, Ly6C^hi^, Ly6G^-^) or polimorphonuclear PMN-MDSC (CD45^+^, CD11b^+^, Ly6C^lo^, Ly6G^+^). Interestingly, we observed a 1.79-fold decrease of M-MDSC (**Fig. 6G and 6I**) and a 3.04-fold decrease of PMN-MDSC (**Fig. 6H and 6I**) in the TME of NPA-shFYN gliomas compared to NPA-NT controls. Further, we analyzed the immunosuppressive function of MDSC isolated from the TME of GEMM tumors. We first characterized MDSC by expression of T cell-immunosuppressive molecules, i.e., as ARG1 and CD80. We observed a significant decrease in the proportion of CD80+ and arginase+ M-MDSCs, and arginase+ PMN-MDSCs in shFYN tumors **(Fig. 5 J, K, L, M)**. To test whether decreased numbers of MDSC in shFYN TME is due to reduced MDSC migration we performed an *in vitro* bone marrow-derived MDSC migration assay **(Fig 6A)**. This experiment showed that shFYN conditioned media reduced M-MDSC migration. Migration of PMN-MDSC was not decreased in the shFYN group (**Fig. 6B).** Finally, we analyzed the functional MDSC-mediated T-cell immune suppressive activity. We observed that PMN-MDSC (GR1^hi^ CD11b+) or M-MDSC (Gr1^low^ CD11b+) from the TME of NPA-NT and NPA-shFYN decreased T-cell proliferation stimulated by SIINFEKL peptide. However, MDSCs isolated from the sh-FYN tumors were significantly less inhibitory. These data suggest that the microenvironment of shFYN glioma tumors reduce the capacity of MDSCs to inhibit T-cell activation.

**Figure 5.**
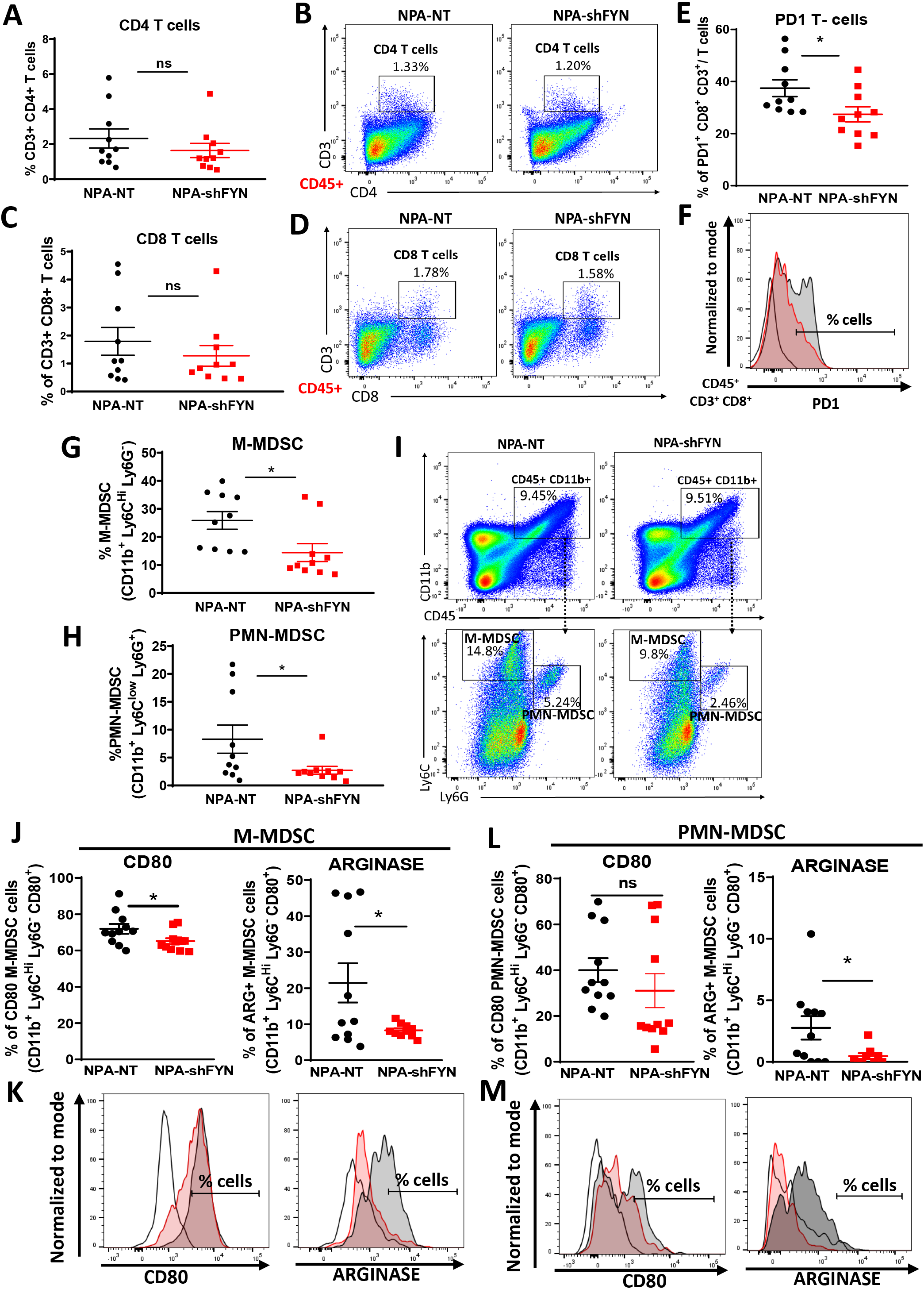
Downregulation of FYN in glioma modulates immune responses through reduction in MDSC expansion. Immune cells within the TME of NPA-NT or NPA-shFYN tumors were analyzed by flow cytometry. **(A-B)** Percentage of CD4-T (CD4^+^, CD3^+^) cells within the CD45^+^ cell population. Representative flow plots for each group are displayed **(C-D)** Percentage of CD8-T cells (CD8^+^, CD3^+^) within the CD45^+^ cell population. Representative flow plots for each group are displayed. **(E-F)** Percentage of PD1+ T cells within the CD8+ CD3+ and CD45^+^ cell population. Representative histogram flow plot for each group are displayed. **(G)** Percentage of monocytic myeloid derived suppressor cells (M-MDSCs: CD11b+, Ly6C^hi^, Ly6G-) within the CD45^+^ cell population. **(H)** Percentage of polymorphonuclear myeloid derived suppressor cells (PMN-MDSCs: CD11b+, Ly6C^lo^, Ly6G+) within the CD45^+^ cell population. **(I)** Representative flow plots of M-MDSC and PMN-MDSC cell analysis. **(J-M)** Percentage of CD80+ and Arginase+ cells within **(J-K)** M-MDSC (CD45+, CD11b+, Ly6C^hi^, Ly6G-) cell population and **(L-M)** PMN-MDSC (CD45+, CD11b+, Ly6C^lo^, Ly6G+) cell population. **(K-M)** Representative histogram of CD80+ and ARGINASE+ cell analysis. Red-shaded: NPA-NT, Grey-shaded: NPA-shFYN, Solid Black-dashed: FMO Control. Each graph indicates individual values and mean ± SEM (n= 10). Data were analyzed using ANOVA test. ns= non-significant; *p < 0.05.

**Figure 6.**
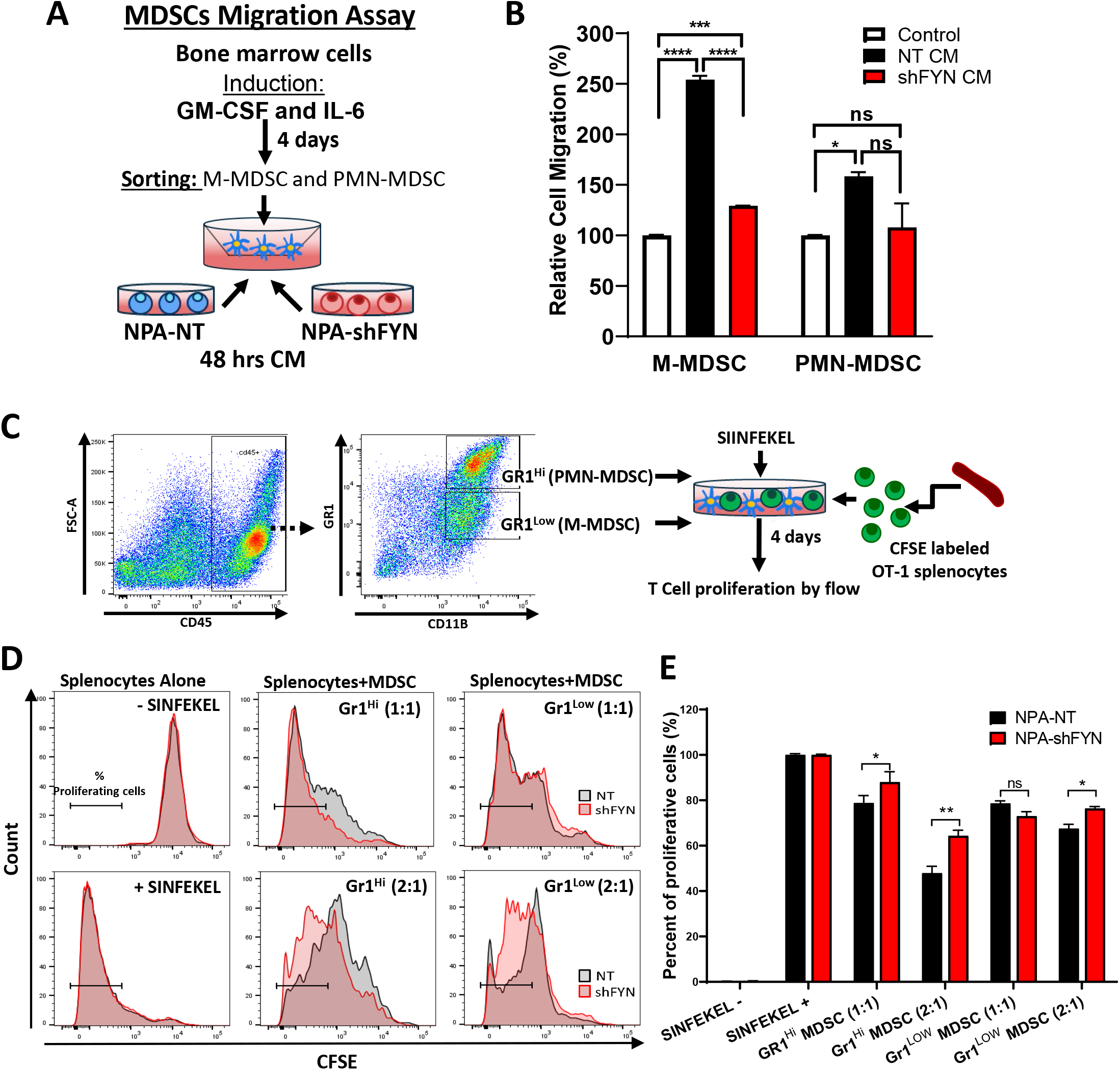
FYN knockdown in glioma decrease MDSC migration potential and immune suppressive activity within the tumor microenvironment. **(A)** Diagram of experimental design of MDSC transwell migration assay. MDSCs derived from bone marrow and induced with IL6 and GM-CSF were seeded on the top of the Transwell and incubated for 15 hours in NT and shFYN conditioned media. The migrated cells were analyzed using CellTiter-Glo^®^. **(B)** MDSC migration assay results. Data is expressed as percentage of migrating cells relative to the control (plain media). Error bars represent ±SEM. Experiment was performed 3 times with 3 replicates per treatment. Statistical significance was determined using One-way ANOVA test, followed by Duncan multiple comparisons. ns: Non-significant, *p<0.05, ***p<0.001, ****p<0.0001. **(C)** Diagram representing the experimental design to analyze the immunosuppressive potential of MDSCs. Gr-1^high^ (PMN-MDSC) and Gr-1^low^ (M-MDSC) were purified from the TME of moribund NPA-NT and NPA-shFYN tumor-bearing mice. They were cultured with CFSE-labeled splenocytes from Rag2/OT-1 transgenic mice. Cells were stimulated with SIINFEKL peptide and proliferation was analyzed 4 days after by flow cytometry. **(D)** Representative flow plots for CFSE stains from splenocytes alone SIINFEKL stimulated and non-stimulated, and the effect of SIINFEKL-induced T cell proliferation in the presence of MDSCs from the TME. Numbers in parentheses indicate the ratio of MDSCs to splenocytes. **(E)** Graph shows T cell proliferation relative to SIINFEKEL+ control. Experiment was repeated 3 times. Tumors from 5 mice per group were pooled together in each experiments to obtain sufficient number MDSCs. Mean ± SEM are indicated. Data were analyzed using one-way ANOVA, followed by Duncan multiple comparisons. ns: non-significant, *p<0.05, **p<0.005.

Besides, we analyzed the expansion and activation status of macrophages. We did not observe a significant increase in the frequency of macrophages (CD45^+^, CD11c^-^, F4/80^+^) (Supplementary **Fig. 8H and I**). The expression MHCII on macrophages in the TME was increased by 1.4-fold in NPA-shFYN gliomas versus NPA-NT controls (**Fig. 8J and K**). It is likely that the lower proportion and suppressive activity of MDSC in shFYN tumors leads to increases of M1 macrophages (MHCII^hi^) and therefore reduced polarization to M2 macrophages. Overall, our data show that downregulating expression of FYN in glioma tumors decreases expansion of MDSC in the TME due to reduced migration potential, decreased expression of CD80 and ARG1, and lower functional immune suppressive activity.

## DISCUSSION

In this study we demonstrate that glioma-cell specific genetic inhibition of FYN tyrosine kinase increases anti-glioma immune-responses, thus significantly delaying tumor progression.

FYN is an effector of the RTK (EGFR, MET, PDGFR) pathway in glioma and other cancers. Downstream of RTK, FYN signals through RAS dependent (via RAS/MEK/ERK) and RAS-independent pathways (via PIK3/AKT, B-catenin, FAK, PXN, STAT3, VAV1 and/or SHC) **(Fig. 1A)**. To activate several downstream molecular pathways, and physiological processes such as cellular proliferation, migration, and cell adhesion^4,17^. RTK are commonly mutated drivers in high grade gliomas; yet their detailed downstream signaling pathways remain incompletely understood^3,4,17–19^.

Molecular analyses of high grade glioma from the Rembrandt, TCGA and Gravendeel databases^7^, and data from Lu *et al*^3^, indicate increased expression of FYN. These human data correlate with our results in mouse models of gliomas, in which increased levels of FYN expression correlate with higher glioma aggressiveness. Further, our molecular gene interaction network analysis highlights FYN as a central network hub, suggesting it might function as a potential regulator of glioma malignancy.

Previous studies have shown that FYN is expressed by both tumor cells and immune cells^20^. In T cells, FYN regulates effector functions and amplifies T-cell antigen receptor (TCR) signaling^4,18,19,21^. However, it has been difficult to establish the role of FYN within glioma in vivo. The function of FYN can be studied genetically (i.e. knockdown) or pharmacologically (i.e. Saracatinib, or Dasatinib inhibitors). Genetic inhibition is highly specific. Pharmacological inhibition, however, is non-specific. Saracatinib or Dasatinib will inhibit all SFK members and will inhibit SFK in all cell types i.e. FYN in the immune cells.

For instance, *in vitro* studies using SFK inhibitors show that FYN promotes cell proliferation and migration in gliomas^3,9,22,23^ *In vivo,* effects of Saracatinib treatment have been mixed, yet unable to determine any specific role of FYN in glioma growth^3,8^. Importantly, Dasatinib, did not extend survival of glioma patients in a Phase II clinical trial^24^. *In vivo* however, there was no effect on animal survival.

As Saracatinib and Dasatinib are non-specific inhibitors of individual SFK member^25^, the genetic inhibition of FYN remains the best option to study its functions in glioma biology. Indeed, genetic downregulation of FYN expression inhibited glioma cells migration and proliferation *in vitro*^3^,^8^,^9^,^26^, yet failed to affect glioma progression in *in vivo* immune suppressive mice^8^.

To address this paradox, we developed a genetic approach to inhibit FYN expression in tumor cells using a GEMM model of glioma in immune competent animals. Our results suggest that shFYN delays tumor initiation and progression *in vivo* by inhibiting the anti-glioma immune response.

We demonstrated the role that FYN plays as central hub in the anti-glioma immune response and tumor progression by inducing tumors in immune-deficient animals. The effect of FYN in delaying tumor growth was abolished in NSG, CD8−/− and CD4−/− immune deficient animals. Both CD8 and CD4 T cells are necessary for increased tumor rejection of sh-FYN gliomas. These studies suggest that FYN plays a crucial role in conveying immune inhibitory messages from within the glioma cells to the immune cells, thereby engineering the local immune response to favor tumor growth. Herein, we uncovered a new cell-non-autonomous mechanism by which FYN knockdown within glioma cells modulates the anti-glioma immune responses.

Analysis of the molecular changes induced by shRNA-FYN strongly suggests that the downregulation of FYN in tumor cells activates the anti-tumor immune response. Meta-analysis of GO for NPF vs NP, NPAF vs NPA and NPDF vs NPD revealed significantly common immune related biological process among all genetic glioma models of FYN-knockdown such as “cellular response to interferon-gamma”, “cellular response to interferon-beta”, “antigen processing and presentation of exogenous peptide antigen via MHC class II”, “cellular response to interleukin-1”, “myeloid dendritic cell differentiation”, “positive regulation of T cell proliferation” and others. Moreover, further functional network analysis found that FYN knockdown tumors display central hubs regulators and over-represented pathways as STAT1, a prominent regulator of the immune system^27^. This module regulates immune functions, IFNγ cellular signaling, JAK-STAT pathways, cell differentiation of Th1, Th2 and Th17, T cell activation and NK cell mediated cytotoxicity^28^,^29^. Although they exhibit the same final phenotype and common biological processes or signaling pathways, the specific mechanisms by which FYN downregulation regulates the anti-glioma immune response would be different for each GEMM of glioma.

Finally, our glioma TME analysis suggests that, FYN downregulation in glioma tumors decreases MDSC expansion and their immune suppressive activity. We and others have previously reported that glioma infiltrating MDSCs play a key role in inhibiting anti-tumor T-cell immune responses, thus promoting tumor progression^16^. The inhibitory immune-microenvironment in glioma is thought to contribute to the ineffectiveness of immunotherapies^30^. Novel therapeutics approaches that reverse the inhibitory microenvironment are essential to counteract these effects, and we propose that the inhibition of FYN within glioma cells could represent such a strategy. shFYN gliomas determined a reduced MDSC migration potential, a decreased immunosuppressive cell phenotype (lower CD80 and ARG1 cells) and lower functional immune suppressive activity. T cell depletion of L-arginine through Arginase causes interference with the CD3ζ chain and proliferative arrest of antigen-activated T cell. Inhibitory CD80 receptors of B7.1 family was implicated in MDSC mediated immune suppression (REF). Glioma microenvironment of FYN knockdown glioma diminished myeloid cell-mediated immunosuppression, leading to increased T cell proliferation and cytotoxicity, decreased T cell exhaustion and M1 to M2 macrophages polarization amplifying anti-glioma immunity.

We propose that inhibition of FYN within glioma cells could be an immune-mediated therapeutic target to restrict MDSC immune suppressive expansion. Since FYN is expressed by both immune cells and tumor cells, therapeutic approaches will need to target FYN specifically in tumor cells. We propose that the combination of tumor cell specific FYN inhibition with other immune-stimulatory treatments, such as immune checkpoint blockade (PDL1-PD1 inhibitors), or Ad-hCMV-TK and Ad-hCMV-Flt3L gene therapy^11,31,32^ are promising avenues that are worth of being explored in future experiments.

## Authorship

### Conception and design

A. Comba and P.R. Lowenstein.

### Development of methodology

A. Comba, P.J. Dunn, A.E. Argento, P. Kadiyala, P.R. Lowenstein.

### Acquisition of data, analysis and Interpretation

A. Comba, P.J. Dunn, A.E. Argento, P. Kadiyala, M. Ventosa, P. Patel, D.B. Zamler, F.J. Nunez, L. Zhao, M.G. Castro, P.R. Lowenstein.

### Manuscript writing

A. Comba, P.J. Dunn, M.G. Castro, P.R. Lowenstein.

### Review, and/or revision of the manuscript

A. Comba, P.J. Dunn, A.E. Argento, P. Kadiyala, M. Ventosa, P. Patel, D.B. Zamler, F.J. Nunez, L. Zhao, M.G. Castro, P.R. Lowenstein.

### Administrative, technical, or material support (i.e., reporting or organizing data, constructing databases)

A. Comba, P.J. Dunn, P.R. Lowenstein.

### Study supervision

M.G. Castro and P.R. Lowenstein.

