## Supplementary Figures for "FYN tyrosine kinase, a downstream target of receptor tyrosine kinases, modulates anti-glioma immune responses"

#### Supplementary Fig. S1

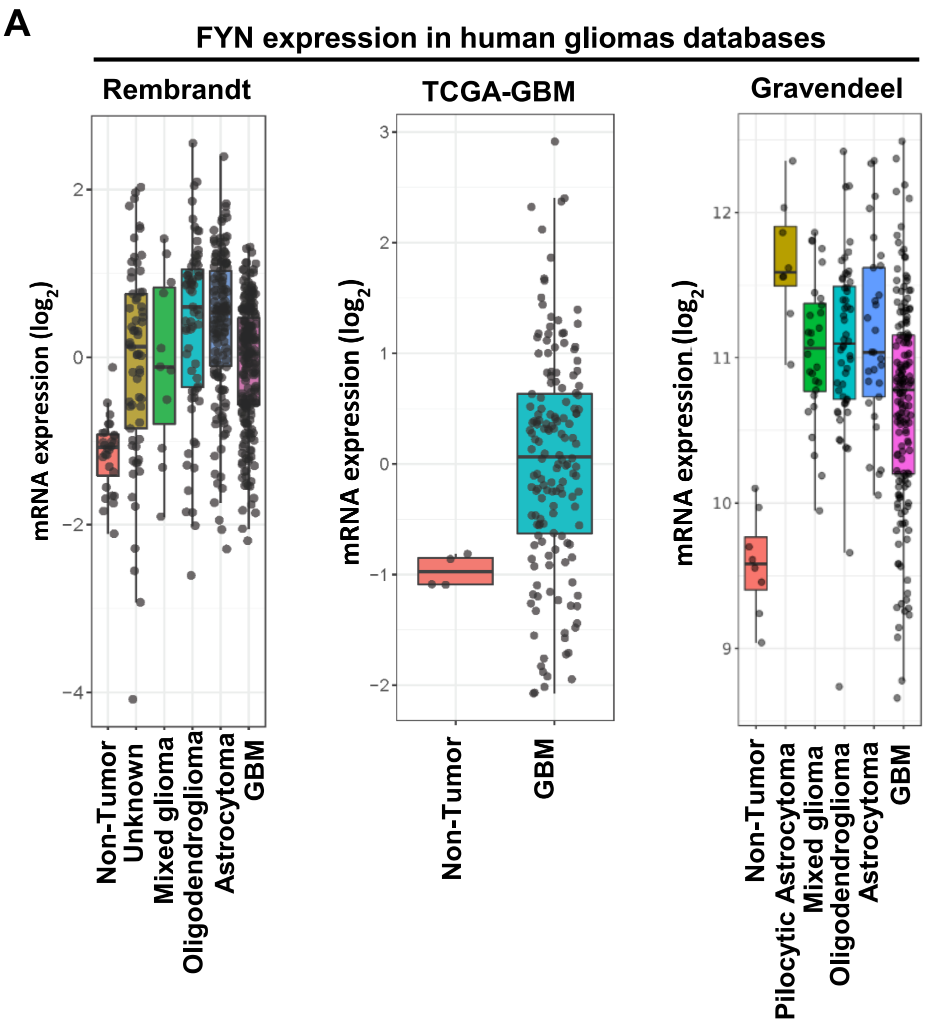


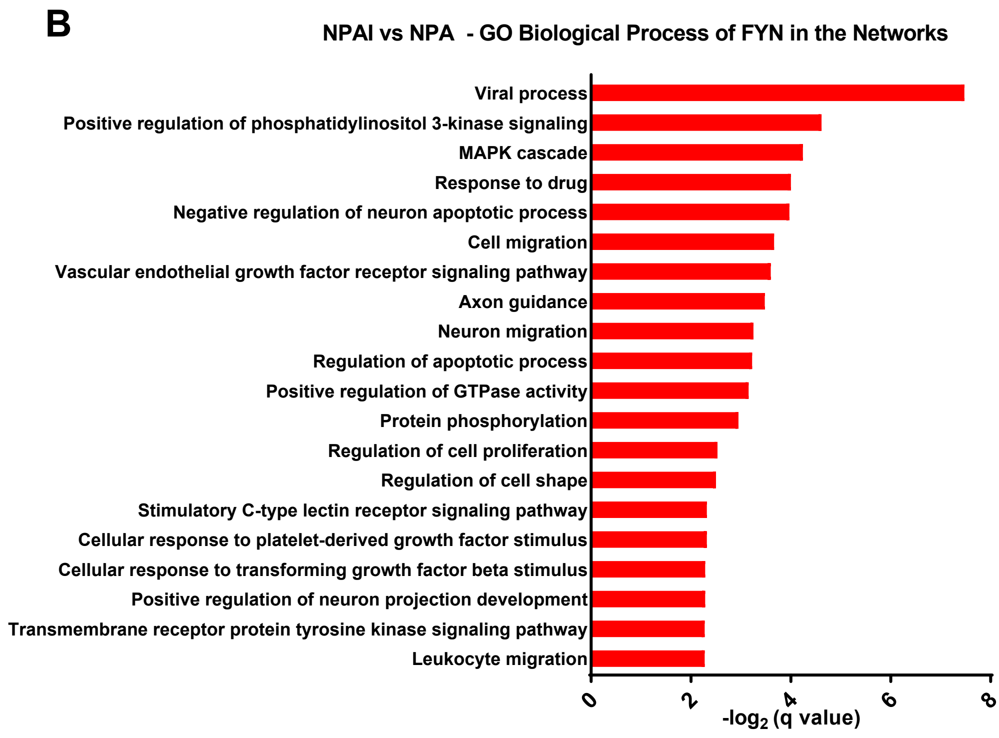

**Supplementary Fig. S2**

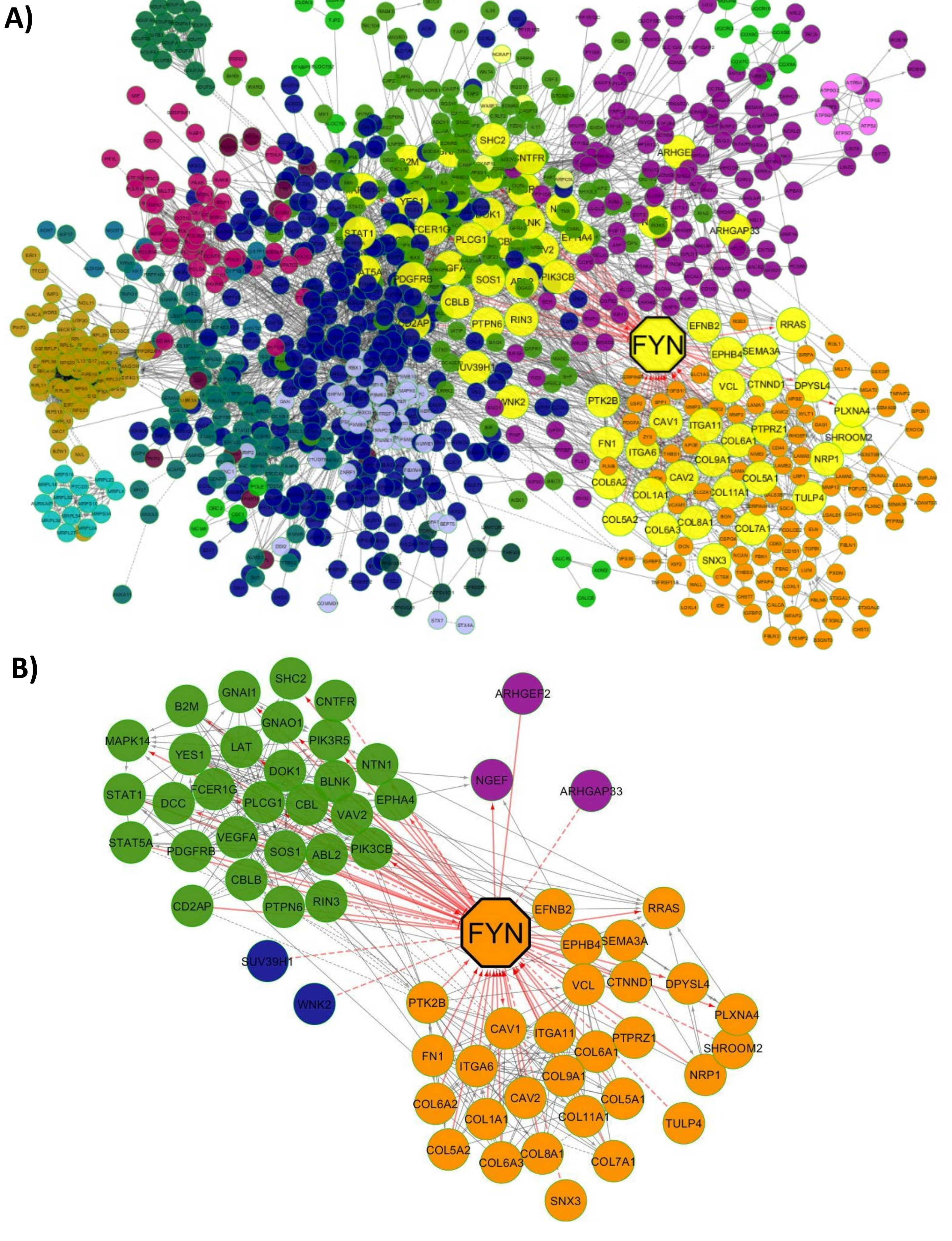

#### Supplementary Fig. S3

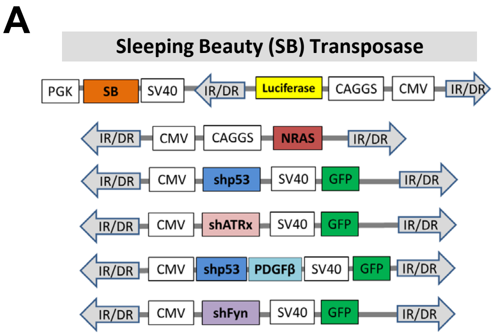


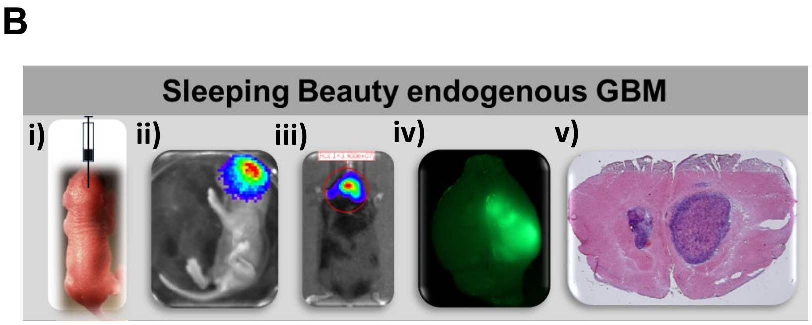


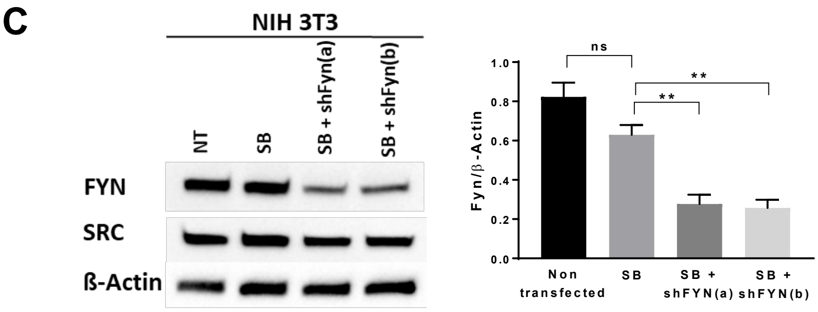


**D**

| Tumor | Genetic Modification |
|---|---|
| NP | N-Ras/shp53 |
| NPF | N-Ras/shp53/shFyn |
| NPA | N-Ras/shp53/shATRx |
| NPAF | N-Ras/shp53/shATRx/shFyn |
| NPD | N-Ras/shp53/PDGFβ |
| NPDF | N-Ras/shp53/PDGFβ/shFyn |

### Supplementary Fig. S4

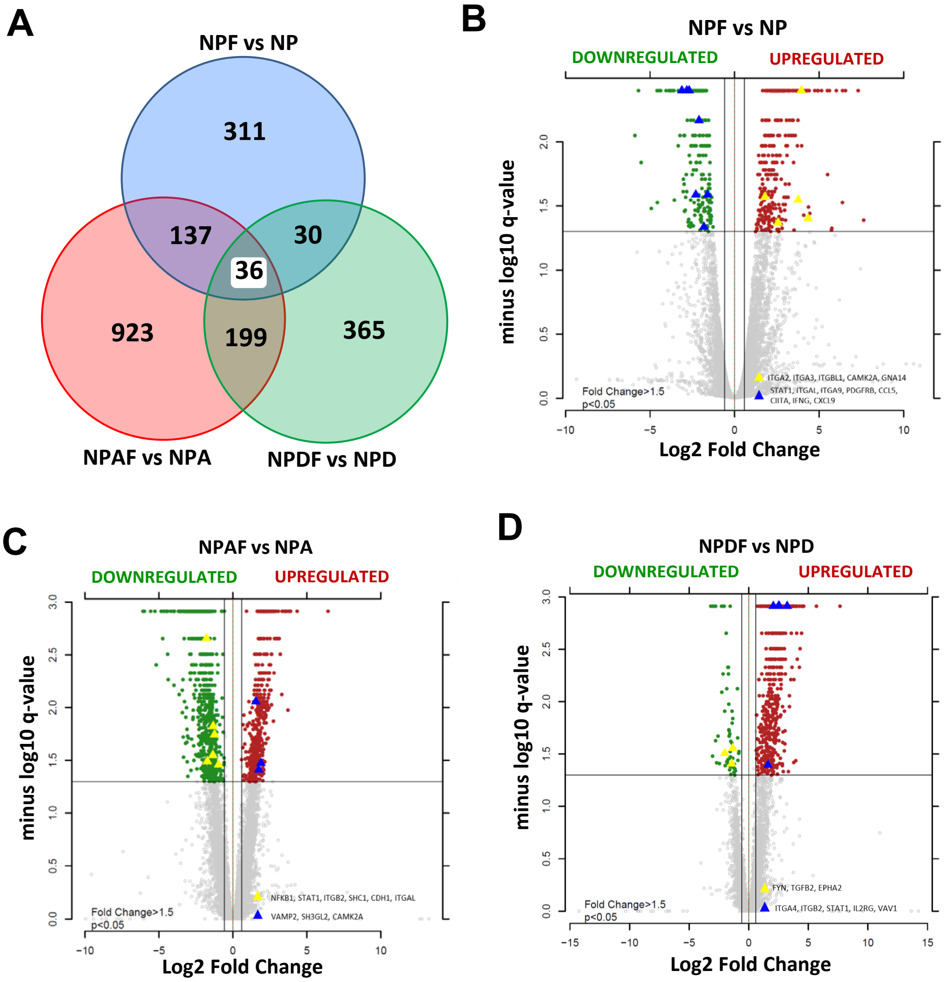

**Supplementary Fig. S5**

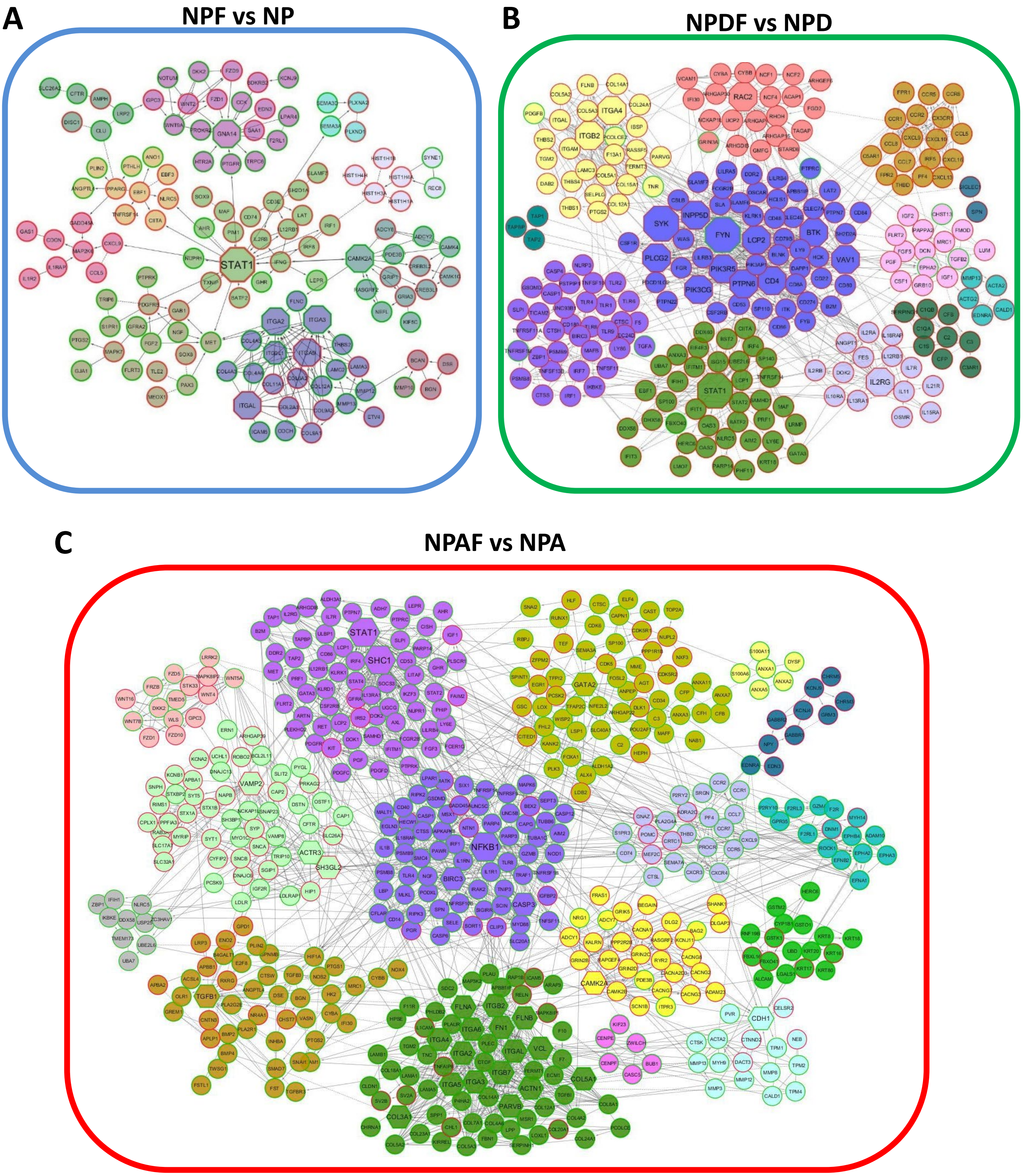

### Supplementary Fig. S6

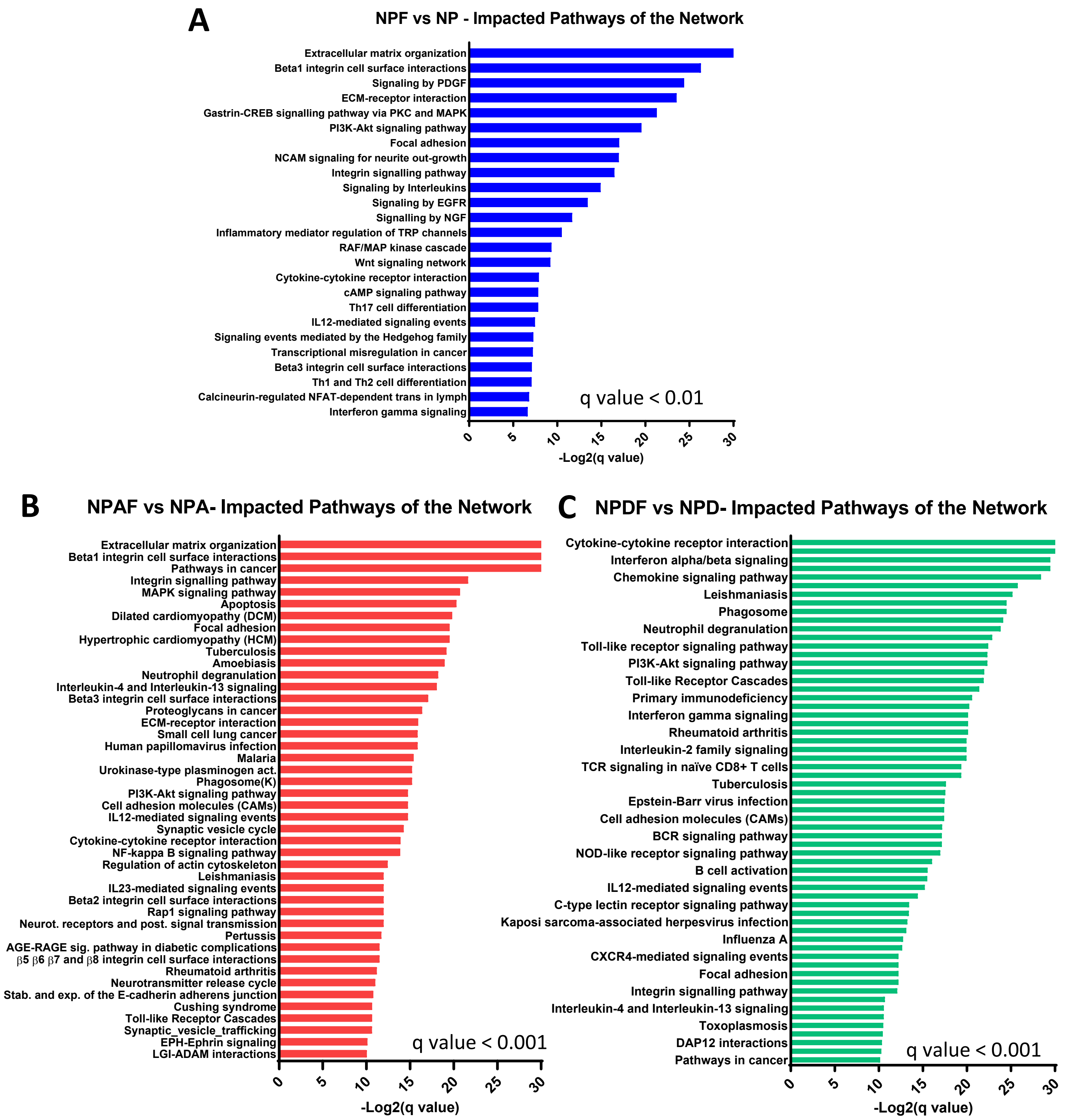

Supplementary Fig. S7

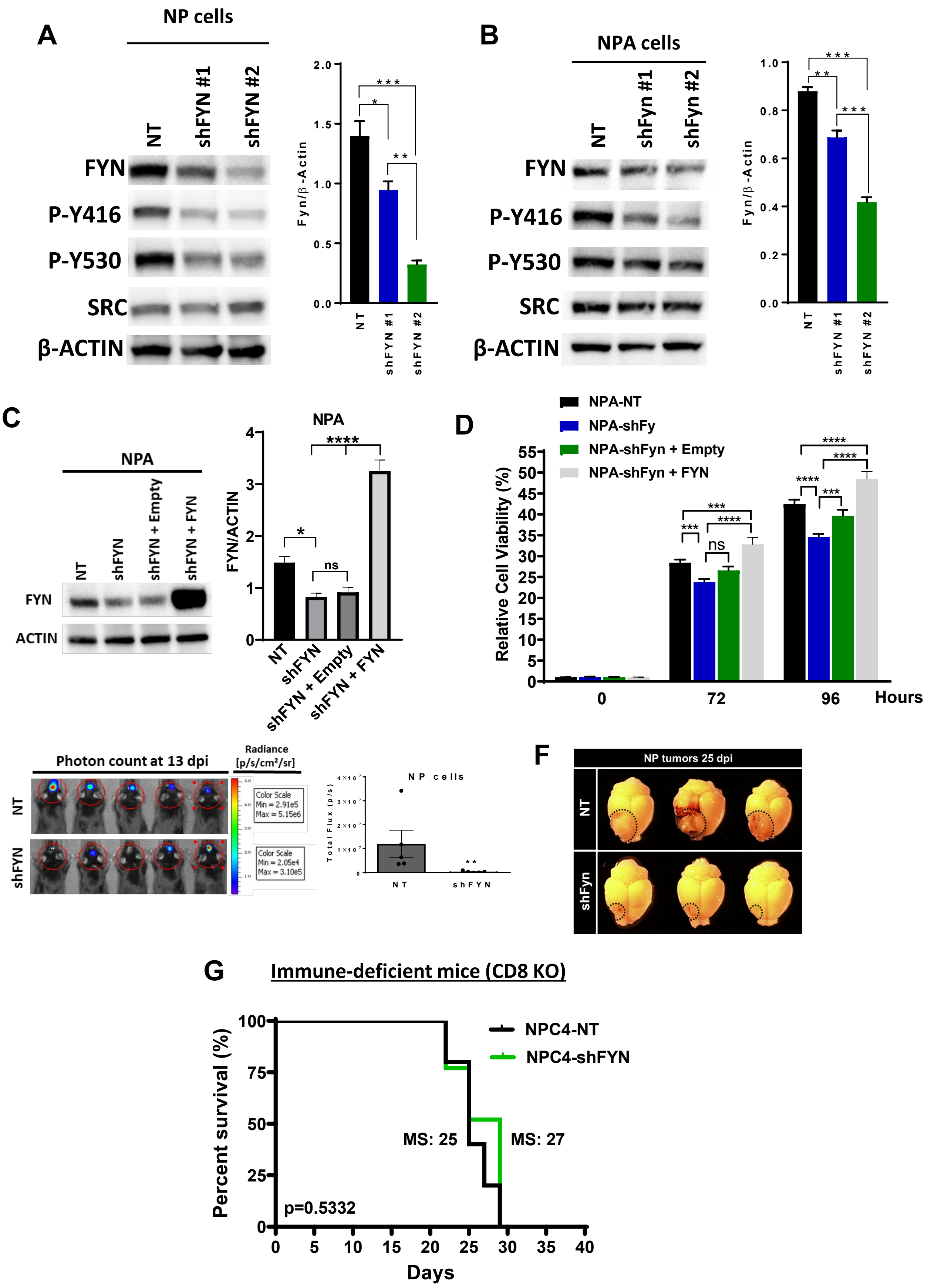

### Supplementary Fig. S8

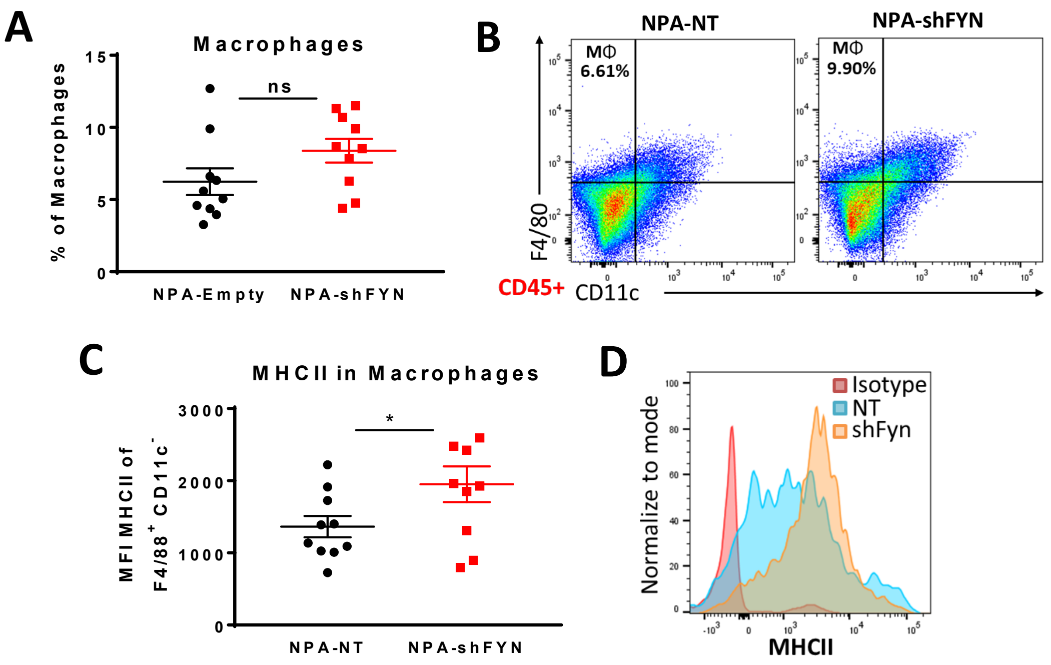
