## Supplementary Information for "FYN tyrosine kinase, a downstream target of receptor tyrosine kinases, modulates anti-glioma immune responses"

### SUPPLEMENTARY DATA

### SUPPLEMANTARY MATERIAL AND METHODS

#### Glioma cell lines and culture conditions

Primary mouse neurospheres: neurospheres were generated from genetically engineered gliomas. Glioma neurospheres exhibit the activation of the RTK/RAS/PI3K pathway, the knockdown of p53 and with or without another specific gene modification as described: NP (N-Ras and shp53), NPA (N-Ras, shp53 and shATRX), and NPAI (N-Ras-shp53-shATRX and mutant $IDH^{R132H}$ expression) [1,2]. All cells used throughout this manuscript: NP, NPF, NPA, NPAF, NPAI, were maintained as glioma neurosphere cultures as described in detail in Koshman C *et al* [1], Nunez *et al* [2]. Their capacity to form tumors upon implantation into mice, or, to differentiate upon plating in the absence of growth factors, justify the description of these cells as glioma stem cells. In this manuscript we now describe them as glioma neurospheres to avoid confusions. Tumors used to generate primary neurospheres were measured using an *in vivo* imaging system to have a bioluminescence reading of $10^7$-$10^8$ photons/s/cm$^2$/sr. Mice were euthanized and brain extracted. The tumor mass was excised through dissection with forceps and scalpel under a fluorescence dissection microscope. The tumor was transferred to 1000 µL of neural stem cell media (NSC) and homogenized mechanically with a plastic pestle. NSC media is formulated as: (DMEM/F12 with L-Glutamine - Gibco®, 11320-033) supplemented with 2% B-27 (Gibco®, 12587-010), 1% N-2 (Gibco®, 17502-048), Penicillin-Streptomycin (Corning, Cellgro®, Corning NY, 30-001-CI), and Normocin™ (InvivoGen, San Diego CA, ant-nr-1). In addition, hFGF and hEGF (Shenendoah Biotech, Warwick PA, 100-26, 100-146) were supplemented twice weekly at 20 ng/ml [1,2]. Tissue was then digested with 1 mL of enzyme-free dissociation reagent, *e.g.*, HyClone™ (GE Healthcare Life Sciences), for 5 min at 37 °C. The homogenized suspension was passed through a 70 µm cell strainer, washed with 10 mL of media, centrifuged at 300 *g* for 4 min, then decanted and resuspended in a fresh 7 mL aliquot of NSC media. The suspended cells were transferred to a T25 flask and placed in a 37 °C incubator (95% air and 5% $CO_2$). After three days, healthy free-floating neurospheres had formed. Cell cultures were maintained at 37 °C with 5% CO2. The media used for the cultures was the NSC media.

**Rescue experiments: generation of stable cell lines for FYN overexpression**

We confirmed the specificity of shFYN results using a non-target shRNA and two additional shRNAs targeting FYN. However, to further address any potential off-target effects we have performed a rescue of the RNAi effects by expressing the FYN cDNA (mutant FYN) in a form refractory to the shFYN. We designed six silent point mutations in the region of the FYN cDNA that is targeted by the shFYN. The shFYN-resistant FYN cDNA (FYN) was synthetized by the Invitrogen GeneArt Gene Synthesis service and cloned into an IRES-containing bicistronic lentiviral vector to allow the simultaneous expression of the mCherry reporter gene (pLVX-mFYN-IRES-mCherry). These plasmid was packaged into lentivirus at the University of Michigan Biomedical Research Vector Core facility and the NPA-shFYN cells were infected with pLVX-FYN-IRES-mCherry virus in the presence of 0.8μg/mL polybrene. Twenty-four hours post-transduction, media was changed and transduced cells were passaged 4 times as needed before been sorted by texas red fluorophore (to enrich stably expressing lines). To confirm the expression of the FYN in the NPA shFYN-FYN-mCherry cells a Western Blotting assay was performed using 1:100 of FYN antibody (Abcam). To further analyze the rescue of the shFYN effects, NPA-NT, NPA-shFYN, NPA-shFYN + Cherry-Empty and shFYN-FYN-Cherry cells were seeded in 96 well plates at a density of 1,000 cells per well. At 1, 2, 3 and 4 days after seeding, cell viability measurements were performed using *CellTiter-Glo*® Luminescent Cell Viability Assay (Promega) according to the manufacturer's protocol.

**Intracranial implantable syngeneic mouse glioma model**

Mice: Immune-competent mice were housed in a pathogen-free, humidity- and temperature-controlled vivarium on a 12:12 hour light: dark cycle with free access to food and water. Intracranial surgeries were performed on 6-8 week old female and male C57BL/6 mice weighing 17-24g in the University of Michigan Unit for Laboratory and Animal Medicine (ULAM).

The strains of mice used to perform *in vivo* studies are as follows: **C57BL6 mice:** C57BL/6J, Jackson stock 000664; H2 *b* haplotype; **NSG mice:** NOD.Cg-Prkdcscid Il2rgtm1Wjl/SzJ, Jackson Stock 005557; H2 $g^7$ haplotype; **CD8 KO mice:** B6.129S2-Cd8atm1Mak/J, Jackson stock 002665; H2 *b* haplotype; **CD4 KO mice:** B6.129S2-Cd4tm1Mak/J, Jackson Stock 002663; H2 *b* haplotype; Thus, control C57BL6, CD8 KO and CD4 KO all display the H2 *b* haplotype. NSG display the H2 $g^7$ haplotype. As NSG mice are devoid of mature B and T cells, dendritic cells and macrophages, serum, and most natural killer (NK) cell cytotoxicity these mice, NSG mice are the most immune-suppressed strain available for studies of human cells and tissues. Thus, they are an excellent model of an immune-deficient animal, and thus perfectly suited for the described experiments. Therefore, the results are explained by the immune components present in each strain, and are not affected by differences in H2.

Intracranial implantation: Cell used for intracranial implantation experiments were: NP-NT and NP-shFYN, and NPA-NT and NPA-shFYN. FYN knockdown stable cell lines were generated using lentiviral vectors expressing shRNA against FYN. WB analysis in NP and NPA cells showed that two shRNAs (shFYN #1 and shFYN#2) decreased FYN protein levels compared with the non-target (NT)-control vector. The shFYN #2 displayed a stronger knockdown for FYN and was chosen for all in vivo studies described further. Prior to implantation, Mice were anesthetized using an intra-peritoneal (i.p.) injection of the anesthetics Ketamine (120 mg/Kg) and Dexmedetomidine (0.5 mg/Kg. Following anesthesia induction, Carprofen (5.0 mg/Kg) was administered subcutaneously. The skull of the mouse was then immobilized in a stereotactic apparatus. An incision was made on the head of the mouse. A burr hole was made using a 0.45 mm drill bit at coordinates corresponding to the striatum (2mm posterior and 1.5mm lateral to the bregma). A 5 µl Hamilton syringe with a removable 33-gauge needle was lowered 3.5 mm ventral into the striatum. Following a two-minute waiting period where the needle was held in place, 1.0 µl of 30.000 neurosphere-derived cells was injected. Each delivery of neurosphere cells was performed over a period of 5 minutes. Following injection, the needle was left in place for an additional minute before being slowly withdrawn from the brain. The incision was sutured with 3-0 nylon sutures. Immediately following surgery, the animals were recovered from anesthesia using Atipamezole via i.p. injection (1.0 mg/Kg) to reverse the Dexmedetomidine. A single subcutaneous injection of Buprenorphine (0.01 mg/Kg subcutaneous) was administered as post-operative pain relief. Sutures were removed 10 days after surgery.

Tumor monitoring: Measurement of tumor development was obtained every week through *in vivo* bioluminescence using the IVIS® Spectrum *In Vivo* Imaging System (Perkin Elmer, USA) until animals showed signs of tumor burden. This methodology allowed us to evaluate tumor evolution. For *in vivo* imaging, mice were anesthetized with oxygen/isoflurane (1.5-2.5% isoflurane). Once anesthetized, 100 µL of luciferin solution was injected intra-peritoneal (ip). Images were taken five minutes after injection of luciferin. To analyze luminescence, Living Image Software Version 4.3.1 (Caliper Life Sciences, Waltham MA) was used. A region of interest (ROI) was defined as a circle over the head, and luminescence intensity was measured using the calibrated unit's photons/s/cm2/sr and the total flux photon/s. For survival analysis, animals were monitored daily for signs of morbidity, including ataxia, impaired mobility, hunched posture, seizures, and scruffed fur. Animals displaying symptoms of morbidity were immediately anesthetized and perfused transcardially with oxygenated Tyrode's solution (20-50 ml). Animals assessed for neuropathology or immunocytochemistry were also perfused with 4% paraformaldehyde as a fixative (100-200 ml). All methods are consistent with the recommendations of the Panel on Euthanasia of the AVMA as documented in their Guidelines for the Euthanasia of Animals. Mouse brains were then removed from the skull and processed for the further analysis.

**Genetically engineered mouse glioma model (GEMM) generation for FYN Knockdown**

Mice: Studies did not discriminate by sex; both male and females were used. All animals were housed in an AAALAC accredited animal facility where they were monitored daily.

Sleeping beauty (SB) transposon system for FYN knockdown glioma murine model: The plasmids used to generate our SB mouse models were completely sequence verified. The plasmid sequences used to generate the tumors were: (i) pT2C-LucPGK-SB100X for transposon & luciferase expression, (ii) pT2-NRASSV12 for NRAS expression, (iii) pT2-shp53-GFP4 for p53 knock-down, (iv) pT2-shATRx-GFP4 for ATRX knock-down, pT2-shp53-PDGFβ-GFP4 for p53 knock-down in combination with PDGFβ ligand overexpression and (vi) pT2-shFYN-GFP4 for FYN knock-down. The pT2CAG-NRASV12 and pT2-shp53-GFP4 plasmids were the generous gift of the Dr. John Ohlfest laboratory (University of Minnesota) [3].

shRNA targeting the FYN gene: To design and cloning of the shRNA targeting the FYN gene (pT2-shFYN-GFP4), we tested two 22 base pair sequences in a 97 base pair hairpin sequence for the mouse FYN gene (shFYN-(1): HP_106460 and shFYN-(2): HP_292369) selected from candidate sequences within the RNAi codex database (http://cancan.cshl.edu/cgi-bin/Codex/Codex.cgi). To confirm the FYN knockdown, NIH/3T3 mouse cells were transfected with the designed shFYN plasmids for 48 hours. Western Blot analysis was performed to analyze the down regulation of the FYN protein in the transfected NIH/3T3 compared to the untransfected control group. FYN protein levels were normalized to β-actin levels.

Neonatal plasmids injections: Female and male postnatal day 1 (P01) wild-type C57BL/6 mice were used for plasmid injections. Plasmid mixture included: (1) SB/Luc, (2) NRAS, (3) shp53, (4) with or without shATRX, (5) with or without PDGFβ and with or without (shFYN). Briefly, plasmids were mixed in mass ratios of [1:2:2]; [1:2:2:2] or [1:2:2:2:2] (20 μg plasmid in a total of 40 μL plasmid mixture) with *in vivo*-jetPEI® (Polyplus Transfection, New York NY, 201-50G) (2.8 μL per 40 μl plasmid mixture) and dextrose (5% total) and maintained at room temperature for at least

15 minutes prior to injection. Neonatal mice (P01) were anesthetized on a barrier over ice for 2 minutes and maintained at a temperature between 2°C and 8°C per ULAM guidelines. Following anesthesia, mice were placed on a neonatal stereotaxic stage cooled to 2-8°C to maintain anesthesia. The lateral ventricle (1.5 mm AP, 0.7 mm lateral, and 1.5 mm deep from lambda) of mice was injected with 0.75 µl plasmid mixture at a rate of 0.5ul/min. Tumor monitoring: To monitor plasmid uptake in neonatal pups, *in vivo* bioluminescence was measured on an IVIS® Spectrum (Perkin Elmer, Waltham MA, 124262) imaging system. One day after plasmid injection, 30 µL of luciferin (30 mg/mL) was injected subcutaneously into each pup.

*Pups* were imaged using the IVIS® Spectrum with the following settings: automatic exposure, large binning, and aperture f =1. Pups without luminescence were euthanized according to the approved protocol. *Adult mice* were monitored daily for signs of morbidity as described before. Tumor growth in adult mice was monitored every 15 days with the IVIS® Spectrum *in vivo* until animals showed signs of tumor burden as described before. This methodology allowed us to evaluate the evolution of the tumors. Survival analysis was performed for each tumor genotype group. Tumors formed within a time-frame dependent on the DNA combination used. Adult mice displaying symptoms of morbidity were transcardially perfused as described before [1,4]. Brain tumors obtained were used for paraformaldehyde-paraffin embedded sections, frozen tumor tissue for RNA-Seq and WB analysis. Live tumor tissue was utilized for neurosphere generation.

**Immunoblotting**

Glioma cells were lysed with RIPA lysis buffer (Sigma Aldrich) with 1x of protease/phosphatase inhibitor cocktail (Pierce®, 78442). 50 µg of protein extract (determined by bicinchoninic acid assay (BCA), Pierce®, 23227) were separated by 4-15% Acrylamide -Polyacrylamide (Bio-Rad, Hercules CA, Mini-PROTEN® TGX™ precast gels) and transferred to nitrocellulose membranes (Bio-Rad, Hercules CA, 1620112) [5]. Membranes were blocked using 5% blocking solution (Bio-rad Laboratories, Hercules, CA, US) diluted in TBS 01% Tween-20 buffer. Membranes were incubated overnight at 4 °C with the following primary antibodies: Rabbit anti-FYN 1:1000, Rabbit anti-SRC 1:1000, Rabbit anti-Phospho-Y416-Src family 1:1000 (Cell Signaling Technology, Inc, Danvers, MA, US), Rabbit Anti-Phospho-Y530 SRC family 1:1000 (Abcam, Cambridge, United Kingdom), and mouse anti-β-actin 1:4000 (Sigma-Aldrich, St. Louis MO, A1978). Then, membranes were incubated with secondary HRP antibody (Dako, Agilent Technologies, Santa Clara CA, goat anti-rabbit 1:4000 (P0448) or rabbit anti-mouse 1:4000 (P0260). Enhanced chemiluminescence reagents were used to detect the signal following the manufacturer's instructions (SuperSignal® West Femto, Thermo Fisher Scientific, 34095). WB quantification was performed using Image J. Reported data is from three individual biological replicates.

**Immunohistochemistry of paraffin embedded brains (IHC-DAB)**

Following perfusion, mouse brains were fixed in 4% PFA (paraformaldehyde) for an additional 48 hours at 4 °C. Brains were then processed and embedded in paraffin at the University of Michigan Microscopy & Image Analysis Core Facility using a Leica ASP 300 paraffin tissue processor/Tissue-Tek paraffin tissue embedding station (Leica, Buffalo Grove IL). Tissue was sectioned using a rotary microtome (Leica) set to 5 μm in the z-direction. Endogenous Peroxidase Quenching was completed through a 0.3% $H_2O_2$ incubation for 5 minutes at room temperature. Heat induced antigen retrieval was performed using 10mM Citric Acid, 0.05% Tween 20, pH 6.0. Tissue permeabilization and blocking was completed using PBS 0.2%Tween- with 5% goat serum for one hour at room temperature. P-Histone H3 (S10) (Cell Signaling Technology, Inc, Danvers, MA, US) primary antibody was incubated overnight at 4 C at a dilution of 1:200. Tissue sections were then incubated with the secondary biotinylated goat anti-rabbit IgG (Vector Laboratories, Burlingame CA, PK-6101) at 1:1000 dilution in PBS with 0.2% Tween-20 overnight at 4 degrees. ABC Avidin-Biotin-COMPLEX Binding reagent (Vectastain Elite ABC kit) and Betazoid DAB Chromogen detection kit (BioCare BDB2004) were used according to the manufacturer's instructions. Images were obtained using bright field from five independent biological replicates

(Olympus BX53 Upright Microscope from Olympus). Ten different fields of each section were selected at random to include heterogeneous tumor areas. Image quantification of positive PH3 cells was performed using Image J, (National Institutes of Health).

**Immunofluorescence of paraffin embedded brains**

After perfusion, mouse brains were fixed in 4% PFA (paraformaldehyde) for an additional 48 hours at 4 °C. Tissue sections were de-paraffinized and hydrated. Heat induced antigen retrieval was performed using 10mM Citric Acid, 0.05% Tween 20, pH 6.0. Permeabilization was performed using 0.5% TritonX-100 in PBS for 30 minutes at room temperature while shaking. Next, endogenous peroxidase was quenched by 3% Hydrogen Peroxide and incubated for 30 minutes at room temperature. Tissue sections were blocked in 10% horse serum and 3% BSA in PBS. FYN primary antibody (NBP1-82685, Novus Biologicals, LLC, USA) was incubated at 4˚C overnight in a humid chamber at a concentration of 1:200 in 3% BSA in PBS. Detection was performed using HRP-conjugated streptavidin was incubated for 60 minutes at room temperature. Alexa Fluor™ 488 Tyramide was incubated for 10 minutes. Nuclei were stained with DAPI (1:1000) in PBS for 5 minutes. Images were acquired with a laser scanning confocal microscope in five independent biological replicates per group (LSM 880, Axio Observer, Zeiss, Germany) with a Plan-Apochromat 20x/0.8 M27 objective. Ten different fields of each section were selected at random to include heterogeneous tumor areas. Images were analyzed and quantified using Image J.

**RNA isolation and RNA-Sequencing**

For RNA-Seq analysis, symptomatic animals were transcardially perfused with cold Tyrode's solution for 15 minutes. Bulk tumors were dissected from the brain using an Olympus szx16 stereo-zoom microscope. Bulk tumor tissue was immediately frozen in liquid nitrogen and kept at -80 ˚C until RNA isolation. For RNA isolation, using the RNeasy Plus Mini Kit, 30 mg of tumor tissue was first mechanically disrupted and then homogenized in 600 ul of RLT Plus lysis buffer with 1% β-mercaptoethanol as recommended. Genomic DNA was removed using the gDNA eliminator spin column. RNA was eluted in 50 ul of RNase-free water. Before library preparation, RNA was assessed for quality using the TapeStation System (Agilent, Santa Clara, CA) using manufacturer's recommended protocols. Samples with RINs (RNA Integrity Numbers) of 8.9 or greater were prepared using the Illumina TruSeq Stranded Total RNA Library Prep kit (Illumina, San Diego, CA) using manufacturer's recommended protocols. 100 ng of total RNA was rRNA-depleted using Ribo-Gone (Takara Bio USA). The rRNA-depleted RNA was then fragmented and copied into first strand cDNA using reverse transcriptase and random primers. The 3 prime ends of the cDNA were adenylated and indexed adapters were ligated. The products were purified and enriched by PCR to create the final cDNA library. Final libraries were checked for quality and quantity by TapeStation and qPCR using Kapa's library quantification kit for Illumina Sequencing platforms (Kapa Biosystems, Wilmington MA) using manufacturer's recommended protocols.

The samples were pooled, clustered on an Illumina cBot and sequenced on the Illumina HiSeq 4000, as paired-end 50 nt reads, according to manufacturer's recommended protocols. Tuxedo Suite software package was used for alignment, differential expression analysis, and post-analysis diagnostics by the University of Michigan Bioinformatics Core. Differentially expressed genes of all tumors were used for gene ontology (GO) analysis. Network analysis was performed using Cytoscape and Reactome App. Detailed analysis are described in Supplementary data.

**RNA-Sequencing data analysis**

Sequencing analyses were performed at University of Michigan Bioinformatics Core as described here. Quality of the raw reads data for each sample was checked using FastQC (version v0.11.3) to identify features of the data that may indicate quality problems (e.g. low quality scores, overrepresented sequences, inappropriate GC content). Reads were aligned to the reference genome including both mRNAs and lncRNAs (UCSC mm10) using TopHat (version 2.0.13) and Bowtie2 (version 2.2.1.). Default parameter settings were used for alignment, with the exception of: "--b2-very-sensitive", which forces the software to spend extra time searching for valid alignments. The raw sequencing data are 125 base paired-end reads. Post alignment QC plots showed high quality reads aligned to the reference sequence. Results were received with an alignment rate of about 50%. FastQC was used for a second round of quality control (post-alignment), to ensure that only high quality data would be input to expression quantitation and differential expression analysis. Cufflinks/CuffDiff (version 2.1.1) was used for expression quantitation, normalization, and differential expression analysis, using UCSC mm10.fa as the reference genome sequence. For this analysis, parameter settings: "--multi-read-correct" was used to adjust expression calculations for reads that map in more than one locus, as well as "--compatible-hits-norm" and "--upper-quartile–norm" for normalization of expression values. Diagnostic plots were generated using the CummeRbund R package. Locally developed scripts were used to format and annotate the differential expression data output from CuffDiff. Genes and transcripts were identified as being differentially expressed based on three criteria: test status = "OK", FDR $\leq$ 0.05, and fold change ≥ ± 1.5. Genes and isoforms were annotated with NCBI Entrez GeneIDs and text descriptions. The volcano plots illustrating total genes and DE genes were produced with the R base package. After RNA-Seq analysis, for Gene Ontology (GO) analysis, all differentially expressed genes of each tumor comparison: NPF vs NP, NPAF vs NPA and NPDF vs NPD were used for analysis in I-Pathway guide. For each Gene Ontology (GO) term as described by Ashburner et al., 2002; and Gene Ontology Consortium, 2004. The number of differentially expressed (DE) genes annotated to the term is compared to the number of DE genes expected just by chance. iPathwayGuide uses an over-representation approach to compute the statistical significance of observing at least the given number of DE genes. The p-value is computed using the hypergeometric distribution as described for pORA in the Pathway Analysis section. To assess the enrichment of GO terms by considering the structure of the gene ontology, this p-value was corrected for multiple comparisons using the elim and weight pruning methods described by Alexa et al., 2006. The elim pruning method iteratively eliminates the genes mapped to a significant GO term from more general (higher level) GO terms, while the weight pruning method assigns weight to each gene annotated to a GO term based on the scores of neighboring GO terms. Multiple comparisons for NPF vs NP, NPAF vs NPA and NPDF vs NPD were performed using meta-analysis from I-pathway guide. We analyzed common DE genes and GOs between all groups.

Network analysis of the differentially expressed genes were performed using Cytoscape and Reactome App. Networks were clustered by Reactome Functional Interaction (FI). Clusters with the same color illustrate network module of highly interacting group of genes in the network. Reactome FI was also used to analyze network and module functions: Pathways Enrichment and GO: Biological process enrichment.

Networks analysis of DE genes form NPA versus NPAI neurospheres were performed using the RNA-seq datasets deposited in NCBI's Gene Expression Omnibus with identifiers GSE94902.

Analysis of the expression levels of FYN in normal tissue and in human glioblastomas were performed using the dataset of Gravendeel, Gill, Grzmil, Kamoun and Rembrandt from Gliovis (http://gliovis.bioinfo.cnio.es) [6]. Correlations analysis in human glioblastomas for FYN and STAT1 mRNA expression levels and the mRNA of the genes related to their pathways were performed using cBioPortal http://www.cbioportal.org [7,8]. The studies were selected from the dataset of TCGA Glioblastoma Multiforme (raw data at the NCI; source mutation data from GDAC Firehose) using all the sequenced patient tumor set.

### Flow cytometry analysis

To study the tumor microenvironment cells by flow cytometry animal were euthanized when they displayed sings of neurological deficits, as required by regulations concerning the treatment of animals. At this time the size of tumors is comparable in all experimental groups, and the whole tumor mass is utilized to extract CD45+ immune cells. Therefore, tumors of comparable sizes were analyzed. In this study there was no attempt to study subregions of these tumors, as the total extent of each tumor was utilized for experimental studies. Mouse brains were dissected and immediately transferred to cold Neurospheres (NS) media and homogenized using a 70 µm nylon strainer and a pestle to obtain a single cells suspension of the tumor. Cells were stained with fixable viability dye-Amcyan 485 (1:1000) for 30 minutes. Nonspecific antibody binding was blocked with CD16/CD32 (1:200) for 10 minutes on ice. Myeloid-derived suppressor cells (MDSCs) were labeled with CD45, CD11b, Ly6C, Ly6G. MDSC were labeled for ARGINASE and CD80 expression and cell population characterization. Macrophages were labeled with CD45, CD11c, F4/80, MHCII. T cells were labeled with CD45, CD3, CD4 and CD8. CD8 T cells were labeled for PD1 expression analysis. All stains were carried out for 30 min at 4 °C with 3× flow buffer washes between live/dead staining, blocking, surface staining, cell fixation, and data acquisition. Flow data were measured using a FACSAria flow cytometer (BD Bioscience) and analyzed using FlowJo version 10 (Treestar).

**T Cell Proliferation Assays**

MDSCs were purified from TME of brain tumor micorenviroment by flow sorting. Tumor mass was dissected out from the brain and homoge-nized using a pestle and a 70 µm nylon strainer to obtain a single cells suspension in RMPI media containing 10% fetal bovine serum. TME-infiltrating immune cells were enriched from the homogenized suspension using a 30%/70% Percoll (GE Life-sciences) density gradient. The purified immune cells were labeled with CD45, CD11b, and Gr-1 antibodies, and MDSCs were purified by flow sorting. MDSC were sorted as CD45+, CD11B+ and GR1$^{hi}$ (PMN-MDSC) and CD45+, CD11B+ and GR1$^{low}$ as performed before in our laboratory (**REF**). Purified PMN-MDSC and M-MDSCs were cultured with CFSE-labeled total splenocytes from Rag2 knockout/transgenic OT-I T cell receptor mice (Jackson laboratory) at different ratios. Cultures were stimulated with 100 nM SIINFEKL peptide (Anaspec) for 4 days. Cells were then stained with anti-mouse CD45, CD3 and CD8 antibodies in flow buffer, and T cell proliferation was analyzed by CFSE dye dilution.

#### *In vitro* MDSC migration assay

We generate in vitro bone marrow-derived MDSC at described by Maringo *et al*. and we performed a transwell migration assay. 1.5 x $10^6$ bone marrow cells were culture for 4 days in presence of IL-6 and GM-CSF mouse recombinant cytokines at a concentration of 40 ng/ml. After, MDSC cells were sorted in two populations, M-MDSC (CD11B+, Ly6Chi, Ly6G-) and PMN-MDSC (CD11B+, Ly6Clow, Ly6G+). Transwell® polycarbonate membrane inserts (Corning Inc.) of 6.5 mm diameter and 8 um pore size were utilized for all assays. A suspension of 50,000 MDSCs cells in 100 ul of RPMI 10% FBS medium was seeded on the top of the transwell insert. The bottom well was filled with 600 ul of conditioned media for 48 hours of NPA-NT and NPA-shFYN cells or fresh media as control. All cells that had migrated through the Transwell membrane were lysed by adding 200 ul of CellTiter-Glo® Luminescent reagent and incubated for 10 minutes at room temperature. The percentage of cells that migrated through the Transwell membrane was measured by luminescence using the Veritas™ Microplate Luminometer (Turner Biosystems, Inc).

#### Statistical Analysis

Statistical analysis performed are described here. In experiments that included one variable, the one-way ANOVA test was used. In experiments with two independent variables, the two-way ANOVA test was employed. A posterior Tukey's multiple comparisons test was used for mean comparisons. Student t-test was used to compare unpaired data from two samples. Kaplan-Meier survival curves were analyzed using the Mantel log-rank test. Linear mixed effects models were also used to compare FYN levels and P-H3-S10 quantification by immunohistochemistry. This model considers that multiple observations per animal are correlated through a random effect. Significance was determined if $p<0.05$. All analyses were conducted using GraphPad Prism (version 6.01) and SAS (version 9.4, SAS Institute, Cary, NC). Statistical tests used are indicated within the figure legends.


### Supplementary Figure legends

**Supplementary Figure S1. Gene expression analysis of FYN in human glioma tumors and mouse and human glioma cells shows a correlation with aggressiveness. (A)**, mRNA expression analysis of FYN in normal brain tissue vs different glioma subtype tissue from Rembrandt, TCGA and Gravendeel database. The data was obtained from the Gliovis (http://gliovis.bioinfo.cnio.es) database. Rembrandt dataset: mRNA expression of FYN in non-tumor brain tissue vs glioma subtypes (Unknown, Mixed glioma, Oligodendroglioma, Astrocytoma and Glioblastoma). Graph shows the $\log_2$ mRNA expression levels of FYN. Statistical significance was given within the corresponding databases (pairwise t-test with Bonferroni correction). Non-Tumor vs Unknown, p-value= $1x10^{-6}$; Non-Tumor vs Mixed glioma, p-value= $8x10^{-3}$; Non-Tumor vs Oligodendroglioma, p-value= $1x10^{-12}$; Non-Tumor vs Astrocytoma, p-value= $5x10^{-15}$; Non-Tumor vs GBM, p-value= $4.3x10^{-8}$. TCGA data: Graph shows the Log2 mRNA expression levels of FYN. Statistical significance was determined using the Pairwise t-test. p-value was determined by Bonferroni correction. Non-Tumor vs GBM, p-value= 5.1E-02. Gravendeel database: Non-Tumor vs Pilocytic Astrocytoma, p-value= 2.7E-08; Non-Tumor vs Mixed glioma, p-value= 1.2E-06; Non-Tumor vs Oligodendroglioma, p-value= 1.2E-7; Non-Tumor vs Astrocytoma, p-value= 2.1E-7; Non-Tumor vs GBM, p-value= 1.1E-04. **(B)** Functional enrichment analysis of the gene ontology (GO) terms of the network obtained from Figures 1d and 1e. The bar graph displays overrepresented GO Biological process that include FYN tyrosine kinase. GO term significance was determined by a cutoff of q-value (FDR) < 0.01. GO terms were plotted against the -Log 2 of the q-value (FDR).

**Supplementary Figure S2. NPA vs NPAI network analysis**

Magnification of network of DE genes in NPA (high malignancy) vs NPAI (low malignancy) mouse glioma neurospheres.

**Supplementary Fig. S3. Genetically engineered mouse glioma models for FYN downregulation.**

**(A)** Schematic representation of the Sleeping Beauty Transposon System (SB) plasmids used to generate gliomas with FYN downregulation. Plasmids include the following DNA sequences: Luciferase, NRAS-GV12, shATRX-GFP, shP53-GFP, shP53-PDGFβ-GFP and shFYN-GFP). **(B)** Representative pictures showing tumor development in the SB genetically engineered mouse model: i) plasmid injection in P1 neonatal mice. ii) bioluminescence imaging of a neonatal mouse 1-day post injection (1 dpi) confirming the efficiency of transduction; iii) bioluminescence showing tumor development in an adult mouse harboring NP-shFYN glioma at 120 dpi; iv) fluorescence image of a mouse brain harboring a NP-shFYN glioma tumor co-expressing GFP at protocol end point; v) H&E staining of a NP-shFYN brain tumor sections at protocol end point. **(C)** Western Blot (WB) analysis shows downregulation of FYN levels. NIH-3T3 mouse cells transfected for 48 hours with the corresponding vectors. NT: Non-transfected cells, SB: cells transfected with the SB transposon vector. SB+shFYN: cells transfected with the SB transposon vector in addition to shFYN(a): HP_106460 or shFYN(b): HP_292369. WB shows that expression levels of Src were unchanged with transfection of the SB+shFYN vectors. β-actin was used as loading control. Bar graphs represent the quantitative analysis of the WB. Values were calculated by normalizing FYN band density to the band density of β-actin. Quantification was performed using ImageJ software. WB quantifications performed in 3 independent experiments. Statistical significance was determined using One-way ANOVA test. $^{**}p < 0.01$, ns: non-significance. Error bars represent ±SEM. **(D)** Genotypes of the different SB tumors generated are indicated in the table.

**Supplementary Fig. S4: Complementary figures for the bioinformatics analysis of the differentially expressed genes of genetically engineered mouse glioma models.**

**(A)** Venn diagram of differentially expressed (DE) genes. Meta-analysis of different genetic glioma models shows individual DE genes and 36 common DE genes shared between all groups. Genes with 0.05 p-value and a log fold change of at least east 0.585 absolute value were considered significant. **(B-D)** Volcano plot displays the DE genes from **(B)** SB NPF vs NP, **(C)**

SB NPAF vs NPA and **(D)** SB NPDF vs NPD mouse GEMM. DE genes were selected based on the fold change (≥ 1.5) and a q-value (FDR corrected p-value) of < 0.05. Upregulated genes (red dots) and downregulated genes (green dots) are shown. Example of genes upregulated (blue triangles) or downregulated (yellow triangles) are indicated in the lower right quadrant. Two-sided Student's t-test was performed to determine statistical significance.

**Supplementary Fig. S5: Network analysis of the differentially expressed genes of genetically engineered mouse glioma models.**

**A-B)** Network analysis of the DE genes comparing **(A)** NPF vs NP, **(B)** SB NPAF vs NPA, and **(D)** NPDF vs NPD mouse genetic glioma models. Large octagonal nodes represent networks hubs. Clusters of the same color represents modules of highly interacting genes. Red border: upregulated genes. Green borders: downregulated genes.

**Supplementary Fig. S6: Pathways analyses of the network of differentially expressed genes of genetically engineered mouse glioma models.**

Functional enrichment analysis of pathways for each network. **(A)** NPF vs NP, **(B)** SB NPAF vs NPA, and **(D)** NPDF vs NPD mouse genetic glioma models. The bar graph shows the overrepresented pathways plotted according to the -log2 FDR q-value (FDR). Cutoff of q-value ≤0.01 for A, and q-value ≤0.0001 for B and C.

**Supplementary Fig. S5. Generation of mouse glioma stable cell lines for FYN knockdown implantable models. (A-B)** Western Blot (WB) analysis of FYN expression in stable mouse glioma cells generated by lentivirus FYN-shRNA infection followed by Puromycin selection. Cells were transfected with the non-target shRNA control vector or one of two different shRNA vectors for FYN (shFYN#1 or shFYN#2). WB analysis illustrates the downregulation of FYN levels in NP cells (a) and NPA cells (b) by shRNA#1 or shRNA#2. Expression of phospho-Y416 and Y-530-

Src family kinases and SRC was determined. β-actin was used as loading control. Bar graphs represent the quantitative analysis of the WB. Values were calculated by normalizing FYN band density to the band density of β-actin. Three independent quantifications were performed using Image J software. Statistical significance was determined using One-way ANOVA test. $*p < 0.05$; $**p < 0.01$; $***p < 0.001$. Error bars represent ±SEM. **(C)** Western Blot illustrate FYN expression in mouse glioma neurospheres after transfection. Cells were transfected with the non-target shRNA control vector, shFYN, shFYN + Empty vector, and shFYN + FYN overexpression. Empty and FYN expression vector was cloned in a pLVX-IRES-mCherry backbone. β-actin was used as loading control. Bar graphs represent the quantitative analysis of the WB. Three independent quantifications were performed using Image J software. Statistical significance was determined using One-way ANOVA test. $*p < 0.05$; $****p < 0.0001$. Error bars represent ±SEM. **(D)** Cell viability assay performed on transfected mouse glioma cells to analyze the response to FYN expression levels. Cell viability was evaluated by CellTiterGlo assay performed at 0 hours, 24, 48, 72 hours and 96 hours. The results are expressed in percent cell viability relative to control, and the statistical significance was determined using two-way ANOVA test. $**p<0.01$, $***p<0.001$, $****p<0.0001$. Error bars represent ±SEM. Experiment was performed 2 times with 6 replicates per treatment condition. **(E)** Animals harboring shFYN tumors showed a significant decrease in bioluminescence signal at 13 days post implantation (dpi). Luminescence intensity was measured using photons/s/cm2/sr and total flux (photon/s). Bar graph represents the luminescence intensity as (photon/s) in five (n=5) animals per group. Error bars represent ±SEM. Statistical significance was determined using a t-test. $**p < 0.01$. **(F)** Representative picture of the tumor size seen from the brain's surface analyzed at 25 dpi of animals harboring NP-NT vs NP-shFYN tumors. Tumors with FYN downregulation displayed decreased tumor compared to the control. **(G)** Kaplan–Meier survival curve for NP-NT vs NP-shFYN glioma in CD8 KO immune-deficient mice. No significant difference was observed in survival. For each implantable model n=5 was used. Statistics were assessed using the log-rank Mantel-Cox test.

**Supplementary Fig. S8. FYN knockdown in glioma increase M1 macrophages polarization within the tumor microenvironment**

**(A-B)** Percentage of macrophages within the $CD45^+$ cell population. Graph shows mean ± SEM (n= 10); ns= non-significant; unpaired t-test. **(C-D)** Geometric mean of MHC $II^+$ expression in macrophages within the $CD45^+$ cell population. Representative flow plots for each group. Graph represent mean ± SEM; n= 10; ns= non-significant; unpaired t-test.
